# Detecting Protein Higher-Order Structural Changes Using Kinase as a Phospho-Labeler

**DOI:** 10.1101/2025.05.07.652599

**Authors:** Asato Maeda, Kosuke Ogata, Yasushi Ishihama

## Abstract

We developed an approach to detect protein conformational changes on a proteome-wide scale by labeling proteins with phosphate groups through *in vitro* kinase reactions. This structural proteomics approach combines residue-specific labeling based on substrate recognition by protein kinases with quantitative phosphoproteomics using phosphopeptide enrichment, enabling proteome-wide and site-specific profiling of protein conformational changes. By performing *in vitro* phosphorylation of myoglobin with and without heat denaturation and of HEK293T cell extracts with and without protease treatment, we demonstrated that differences in substrate protein structure are reflected in *in vitro* phosphorylation efficiency. Moreover, we successfully identified protein conformational changes upon RNA digestion in non-denatured cell extract proteome samples on a proteome-wide scale with high sensitivity by comparing phosphorylation efficiencies. This approach enables residue-specific readout of structural dynamics within the intracellular proteome.

**Graphical abstract:** 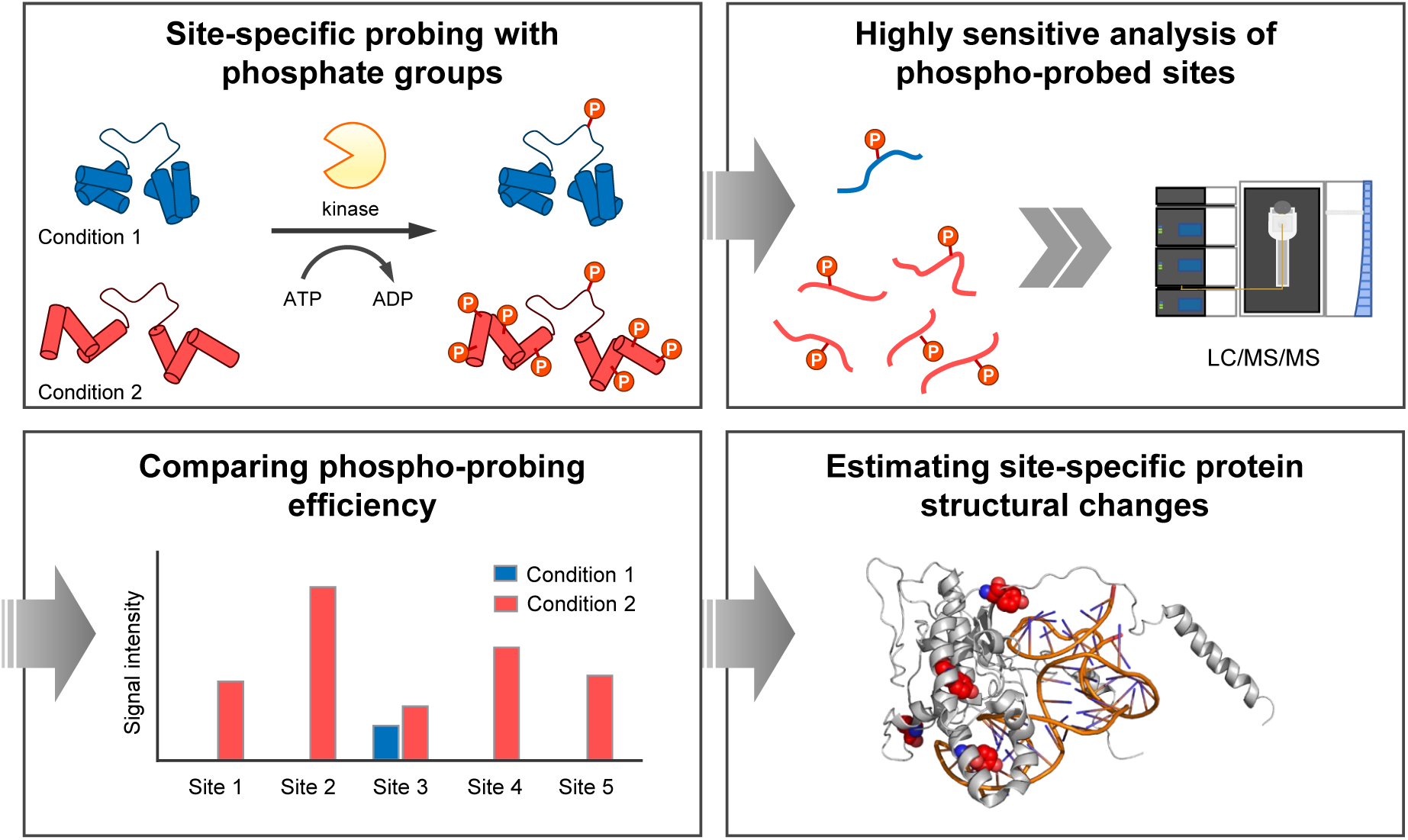

**Motivation:** Mass spectrometry (MS)-based structural analysis of proteins often involves labeling of exposed amino acid residues, followed by digestion and identification of labeled peptides by MS. However, the heterogeneity of labeled peptides and the complexity of biological samples hinder proteome-wide analysis of protein higher-order structures. To address this issue, we devised an approach that utilizes site-specific labeling with phosphate groups, taking advantage of the substrate recognition provided by protein kinases. The phosphate groups serve as enrichment handles, enabling selective extraction and highly sensitive analysis of labeled sites.

**Highlights:**

- Protein labeling with phosphate groups using *in vitro* kinase reaction
- Quantitative MS-based readout of *in vitro* phosphorylation efficiency
- *In vitro* phosphorylation efficiency reflects protein conformational changes
- Proteome-wide and site-specific identification of protein conformational changes

## Introduction

The dynamic structure of proteins is intimately linked to their function^1,2^. In the crowded intracellular environment, protein molecules undergo continuous conformational changes through interactions with other proteins and biomolecules, as well as through post-translational modifications and degradation. These conformational dynamics often drive signal transduction and regulate a wide range of cellular processes by transmitting signaling information to downstream proteins^3,4^. However, such intermolecular communication is absent in purified protein systems, which are commonly employed to study protein structural properties using conventional biophysical methods. To fully elucidate the molecular mechanisms underlying physiological and pathological phenotypes, it is necessary to investigate the structural landscape of the intracellular proteome^5^. Therefore, a comprehensive analysis of protein higher-order structural dynamics—referred to as structural proteomics—is essential for understanding the molecular basis of biological phenomena.

Protein structural analysis techniques combined with mass spectrometry (MS) are increasingly being employed to investigate higher-order structural dynamics at the proteome-wide level^6,7^. These techniques can be broadly categorized into labeling-free and labeling-based approaches. Labeling-free approaches, such as thermal proteome profiling (TPP), detect changes in the physicochemical properties of proteins, enabling comprehensive identification of proteins in which conformational changes are induced by ligand binding, nucleotide binding, and protein-protein interactions^8,9^. However, since these methods detect physicochemical changes for the entire protein, they do not provide site- or domain-specific information on structural changes. Although quantification at the peptide level can elucidate how specific domains affect proteoform stability^10^, differences in the stability of individual domains within the same molecule cannot be resolved. In contrast, labeling-based approaches involve modifying exposed amino acid residues through chemical or enzymatic reactions, followed by identification of the labeled sites (i.e., labeled peptides). These approaches afford high-resolution structural insights by enabling site-specific detection of protein conformational changes. Representative techniques such as hydrogen-deuterium exchange mass spectrometry (HDX-MS)^11,12^, hydroxyl radical protein footprinting (HRPF)^13^, and carbene footprinting^14^ are widely used to identify protein–RNA interaction sites and ligand-binding sites by chemically labeling solvent-accessible residues. However, the heterogeneity of labeled peptides generated by amino-acid-nonspecific labeling reactions complicates mass spectrometric detection^15,16^, limiting the utility of this approach in complex biological samples. To address this limitation, several strategies have been developed to reduce the heterogeneity of labeled peptides and enable site-specific profiling of protein structural dynamics on a proteome-wide scale. One such strategy, limited proteolysis coupled with mass spectrometry (LiP-MS)^17^, monitors proteome-wide conformational changes by analyzing peptides generated by proteases with broad specificity. LiP-MS has recently been applied in a wide range of studies, including the analysis of protein–protein interactions and the identification of drug target sites^18^. However, accurate identification of LiP-derived peptides remains challenging due to the nonspecific cleavage and the increased complexity associated with two-step digestion. While a recent study employing one-step digestion achieved high sensitivity in detecting structural changes, the resulting structural resolution was limited to the peptide level^19^. An alternative strategy involves the use of dimethyl labeling reagents or amino-acid-specific labeling reagents^20–23^. These approaches enable proteome-wide monitoring of structural changes with residue-level resolution. Nevertheless, the wide dynamic range of intracellular protein concentrations demands extremely high residue specificity from chemical reagents, which is difficult to achieve. Enrichment strategies have also been developed to enable sensitive detection of low-abundance labeled peptides. Several reagents incorporating enrichment tags have been reported for use in proteome-wide structural analysis^24,25^. Unfortunately, biotin, the most widely used enrichment handle, binds strongly to streptavidin, making elution difficult^26–28^. Furthermore, the fragmentation patterns of biotinylated peptides are often complex and poorly understood, making site-specific identification challenging^29,30^. Overall, residue-specific labeling and the sensitive detection of labeled peptides continue to be major bottlenecks in system-wide protein structural analysis.

Phosphorylation is a post-translational modification that is introduced onto specific residues in proteins. Phosphopeptides derived from proteins with this modification can be readily and efficiently enriched using well-established metal affinity chromatographic methods^31^. The addition of phosphate groups is regulated by protein kinases, which catalyze the transfer of phosphate groups from ATP to amino acid residues of substrate proteins. Kinases are usually categorized into serine/threonine kinases and tyrosine kinases. Through the substrate recognition mechanisms provided by each kinase, these residues are phosphorylated with high specificity^32^. Despite this specifically regulated process, phosphorylation reactions typically occur within milliseconds^33^. Moreover, purified active kinases are commercially available and have been widely utilized in *in vitro* kinase assays combined with mass spectrometry to investigate kinase-substrate interactions. To date, *in vitro* kinase assays have been employed to profile the sequence specificity of kinases^34,35^, to identify direct substrates and extract consensus motifs surrounding phosphorylation sites^36,37^, to estimate phosphorylation stoichiometry^38^, and to enhance the detection sensitivity of endogenous phosphorylation sites^39,40^.

Here, we aimed to develop a method to profile protein conformational changes on a proteome-wide scale by utilizing kinase as a phospho-labeler. This approach, referred to as phospho-probing, detects protein conformational changes in cellular or cell-extracted proteins with phosphate-modifiable residues by measuring changes in the efficiency of *in vitro* kinase reactions in the cell lysate. In this study, we employed the tyrosine kinase SRC, that has been shown to phosphorylate a broad range of tyrosine residues *in vitro*^36,39^, to perform phospho-probing of substrate proteins in distinct conformational states. Our results demonstrate that *in vitro* phospho-probing can detect conformational changes in heat-denatured myoglobin and can differentiate protein conformational states in cell extracts. Furthermore, we applied this method to analyze protein higher-order conformational changes due to the loss of RNA, and we successfully identified residue-specific conformational changes associated with RNA dissociation.

## Results and Discussion

### Phospho-probing workflow

We present a strategy, termed *phospho-probing*, to analyze higher-order protein structures using an *in vitro* kinase assay. In the first step, proteins and protein complexes extracted under non-denaturing conditions undergo phospho-labeling **(Fig 1)**. Phosphorylation is carried out through a brief *in vitro* kinase reaction using a recombinant kinase, which adds phosphate groups specifically to phosphoacceptor residues located within sequences recognized by the kinase. The proteins are then digested, and the resulting phospho-labeled peptides are selectively enriched from the peptide mixture via phosphopeptide enrichment. The enriched samples are analyzed by nanoLC/MS/MS to identify and quantify the phosphorylated peptides. A control sample incubated without recombinant kinase is used to account for any remaining endogenous kinase activity or endogenous phosphorylation. Comparison with this control allows normalization of phosphorylation introduced by endogenous kinases and enables definition of the phosphorylation level specifically attributed to the recombinant kinase (phospho-probing efficiency). By comparing phospho-probing efficiencies under two or more conditions, conformational changes induced by structural perturbations can be detected.

**Figure 1.**
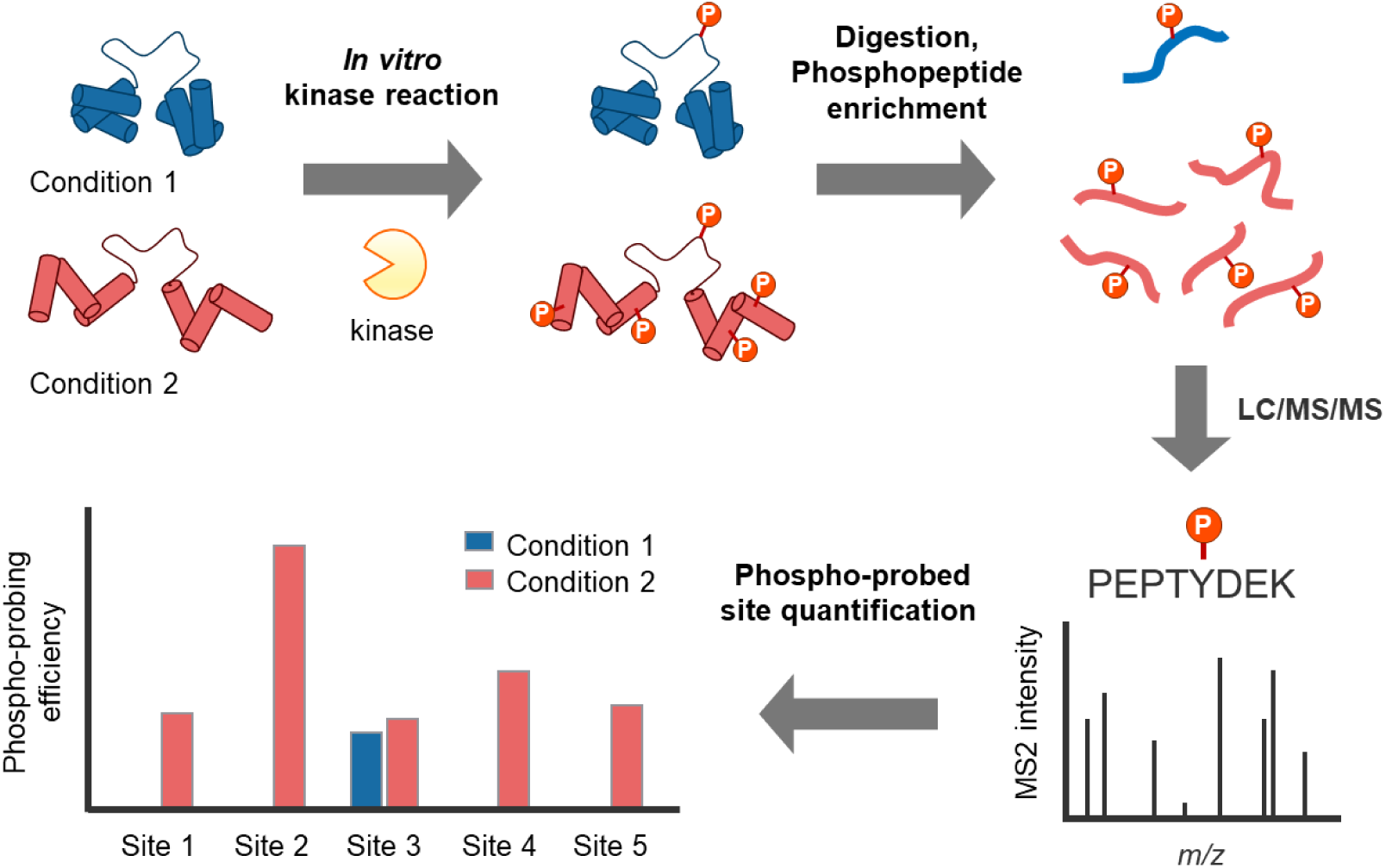
Phospho-probing workflow. In the first step, the native (non-denatured) protein is phosphorylated through a brief *in vitro* kinase reaction using a recombinant kinase. The reaction is terminated by heat treatment of the sample in the presence of a surfactant. Following trypsin digestion, the resulting phosphopeptides are enriched by hydroxy acid-modified metal oxide chromatography with titanium dioxide (TiO₂). The samples are then quantitatively analyzed by LC/MS/MS to detect residue-specific sites of protein conformational change, based on the signal intensity ratios of the identified phosphorylation sites.

In this study, tyrosine residues were selected as targets for phospho-probing, and recombinant SRC kinase, which exhibits high catalytic activity among tyrosine kinases^36^, was used to perform phospho-probing reactions on substrates in various higher-order conformational states. Tyrosine residues are distributed across both hydrophilic and hydrophobic regions of proteins^41^. Therefore, their labeling efficiency is expected to vary significantly depending on the protein’s conformational state at each site. In addition, tyrosine kinases generally exhibit lower substrate sequence specificity than serine/threonine kinases^42^, and many tyrosine residues can be phosphorylated *in vitro*^36,39^. Moreover, the overall level of endogenous tyrosine phosphorylation in mammalian cells is extremely low compared to serine/threonine phosphorylation^43^, which minimizes the influence of endogenous phosphorylation on the analysis of phospho-probed sites.

### Phospho-probing detects higher-order structural changes in heat-denatured myoglobin

We first investigated the feasibility of detecting protein conformational changes by phospho-probing using denatured purified proteins. The experiment was conducted using apomyoglobin from horse, a commonly used model for studying protein conformational changes upon thermal denaturation^44,45^. Since myoglobin contains two tyrosine residues, we first evaluated whether SRC kinase phosphorylates these residues. *In vitro* kinase reactions with SRC were performed on apomyoglobin in the non-denatured state. After trypsin digestion, peptides containing phosphorylated tyrosine residues were quantified in both SRC-treated and SRC-untreated samples. The results showed that Y147 was phosphorylated by SRC, whereas Y104 was not **(Fig S1a)**. We then examined whether the phospho-probing efficiency of Y147 reflects the higher-order structure of the protein. Apomyoglobin was heat-treated at temperatures ranging from 37 °C to 72 °C for 15 minutes. Secondary structure analysis by circular dichroism (CD) spectroscopy or *in vitro* kinase reaction was subsequently performed at 37 °C **(Fig 2a)**. Samples were placed on ice until CD measurements or until kinase reaction was initiated. CD spectra of heat-treated myoglobin showed a decrease in α-helical content, based on ellipticity at 222 nm, confirming that the protein was denatured **(Fig S1b)**. The melting temperature of apomyoglobin was consistent with values found in previous studies^46^. Importantly, the phospho-probing efficiency of Y147 in heat-treated myoglobin increased as the α-helical content of myoglobin decreased **(Fig 2b, Fig S1c)**. These results indicate that phospho-probing can detect changes in the higher-order structure of the protein.

**Figure 2.**
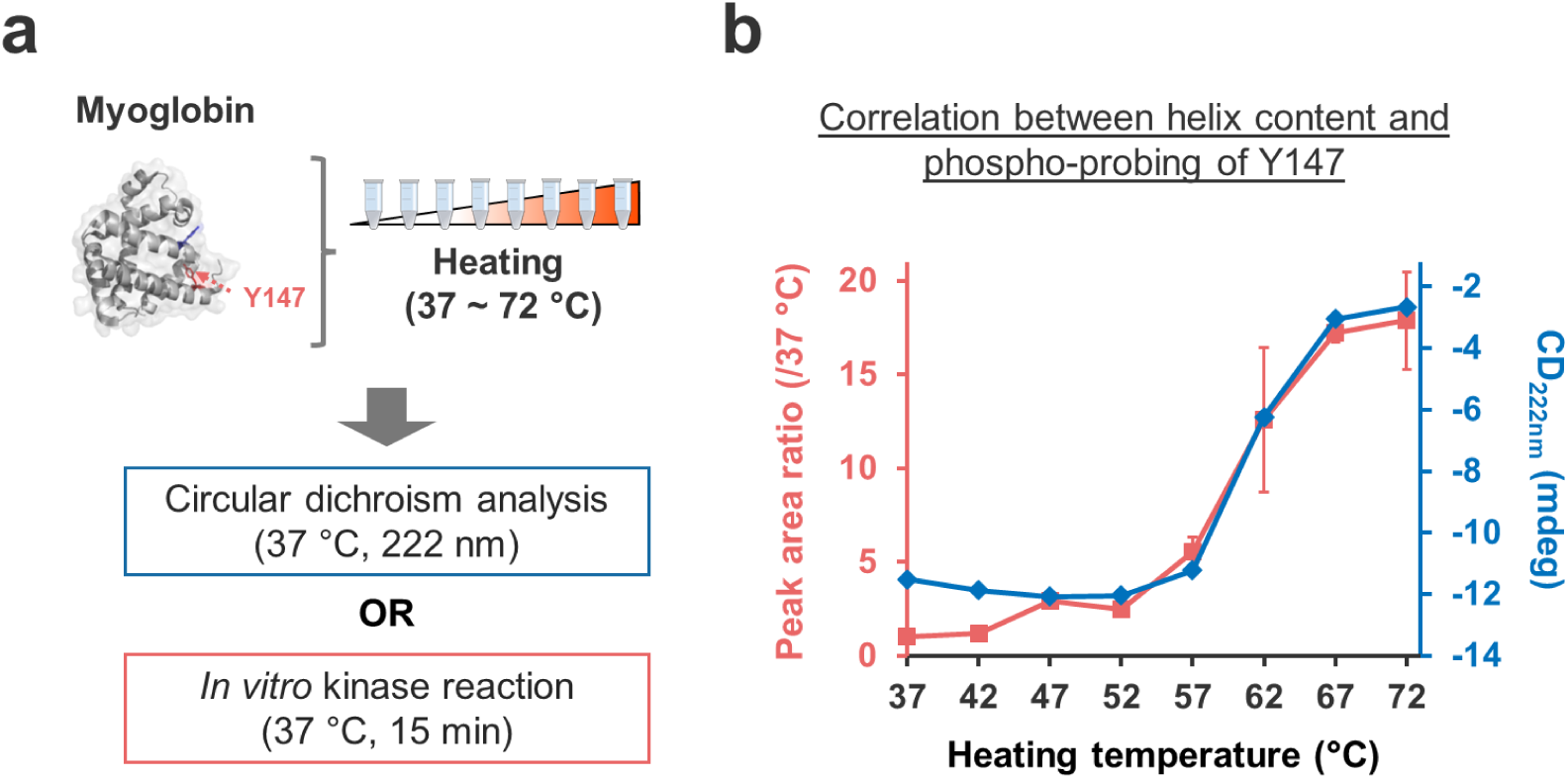
Phospho-probing detects structural changes in heat-denatured apomyoglobin. (a) Workflow of an experiment demonstrating the phospho-probing strategy with apomyoglobin as a model protein. Apomyoglobin from horse was dissolved in reaction buffer and then thermally denatured by heat treatment at temperatures ranging from 37 °C to 72 °C. Secondary structure analysis by circular dichroism (CD) spectroscopy and *in vitro* kinase reaction were performed at 37 °C. The red dotted arrow indicates residue Y147. The structural image was generated based on PDB file 1AZI. (b) Comparison of the secondary structure profile of heat-denatured apomyoglobin and its phospho-probing efficiency. The α-helical content of apomyoglobin was assessed by monitoring ellipticity at 222 nm. Phospho-probing efficiency was calculated as the ratio of the peak area of phosphorylated Y147-containing peptides in each heat-treated sample to that of the control sample (heat-treated at 37 °C). The values represent the average of at least two out of three independent experiments. Error bars indicate the standard deviation.

### Phospho-probing detects the disruption of proteome high-order structure by protease digestion

Next, to evaluate whether phospho-probing can distinguish differences in the higher-order structure of substrate proteins in biological samples, cell lysates were subjected to phospho-probing. Proteins were extracted under non-denaturing conditions by suspending HEK293T cells in surfactant-free buffer under ice-cold conditions, followed by mechanical shearing. The protein extracts were used as a substrate pool for *in vitro* kinase reactions with SRC. To evaluate phospho-probing of substrates with disrupted higher-order structures, *in vitro* kinase reactions were also performed following trypsin digestion of the extracted proteins **(Fig. 3a)**. We identified 440 tyrosine phosphorylation sites in the non-denatured samples and 678 sites in the trypsin-digested samples, in at least two out of three replicate experiments **(Fig. 3b)**. Most tyrosine phosphorylation sites identified across the three replicates overlapped **(Fig S2a)**. However, only 247 phosphorylation sites (56 %) were common to both the non-denatured and trypsin-digested samples, with 193 and 431 sites uniquely identified in the non-denatured sample and trypsin-digested samples, respectively. These results suggest that phospho-probing profiles differ substantially depending on the higher-order structure of the substrate proteins.

**Figure 3.**
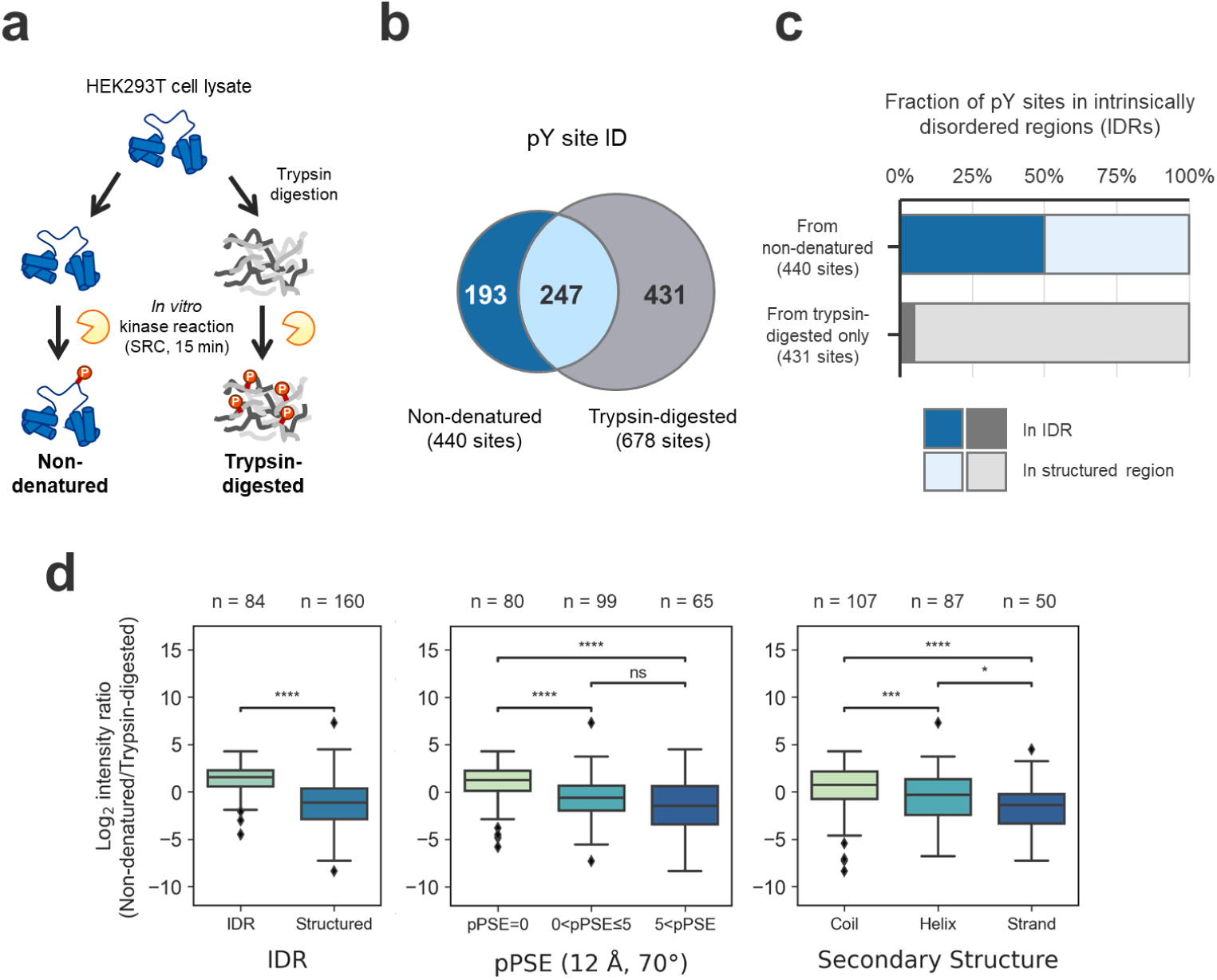
Phospho-probing of cell lysates in the non-denatured state. (a) Experimental design for phospho-probing of non-denatured cell lysates. To evaluate labeling efficiency on substrates with disrupted structural integrity, trypsin digests were prepared from extracted proteins, followed by phospho-probing and phosphopeptide enrichment. Samples were analyzed by nanoLC/MS/MS. Phospho-probing reactions were performed in triplicate. (b) Venn diagram showing the overlap of tyrosine phosphorylation sites (pY sites) identified in non-denatured and trypsin-digested samples. Numbers indicate tyrosine phosphorylation sites detected in at least two out of three technical replicates. (c) Proportion of tyrosine phospho-probed sites located in intrinsically disordered regions (IDRs), as predicted by StructureMap^47^, among all identified tyrosine phospho-probed sites. Tyrosine residues from proteins not available in the AlphaFold database were excluded from the analysis. (d) Boxplot showing the comparison of phospho-probing efficiency of non-denatured lysate according to predicted disorder, solvent accessibility (pPSE at radius = 12 Å and angle = 70°), and predicted secondary structure. The box itself spans the interquartile range. The whiskers represent 5 % and 95 % quantiles. The thick horizontal line in each box indicates the median. Asterisks indicate results of Mann-Whitney U test (ns: p>0.05, *: p<0.05, ***: p<0.001, ****: p<0.0001). Tyrosine residues of proteins for which no predicted structure was available in the AlphaFold database were excluded from the analysis.

We next investigated why phospho-probing profiles vary widely depending on the higher-order structure of the substrate. To explore the relationship between phospho-probing sites and protein higher-order structure on a proteome-wide scale, we mapped the identified phospho-probing sites to predicted protein structures obtained from AlphaFold2^48^. This allowed us to extract comprehensive structural information for the labeled residues. Specifically, the StructureMap python package was used to retrieve residue-level structural predictions from the AlphaFold Protein Structure Database, and intrinsically disordered regions (IDRs) were predicted for each residue^47^. As a result, 859 of the 871 identified tyrosine phosphorylation sites were successfully mapped to predicted structures corresponding to 665 proteins. The remaining sites could not be mapped because the AlphaFold database did not provide structural predictions for proteins with atypical sequences, such as those longer than 2700 amino acids or containing non-canonical residues. Among the 440 sites identified in the non-denatured samples, 434 were mapped, and 50 % of these tyrosine residues were located in IDRs. In contrast, among the 431 sites uniquely identified in the trypsin-digested samples, 425 were mapped, with more than 95 % of the tyrosine residues located in structured regions **(Fig 3c, Fig S2b)**, indicating that phospho-probing profiles reflect the increased accessibility of tyrosine residues upon disruption of protein higher-order structures.

Interestingly, we also observed 193 tyrosine residues that were phospho-probed in the non-denatured substrate, but not in the trypsin-digested sample. These differences may be explained by multiple factors: trypsin digestion may remove amino acid sequences required for kinase recognition, phosphorylation in the non-denatured state may inhibit trypsin cleavage near phosphorylation sites, or distal domains essential for kinase access may be lost during digestion. While all three hypotheses are plausible, we specifically examined the first one, because this hypothesis is supported by the observation that the distribution of lysine and arginine residues around tyrosines that were no longer phosphorylated in the tryptic digest differed significantly from those that were phosphorylated in the tryptic digest (p = 0.024, Fligner–Killeen test) **(Fig S2c)**, suggesting that some phospho-probed sites in the non-denatured samples were affected by trypsin digestion. To further characterize the phospho-probed sites uniquely identified in non-denatured samples, we compared them with phospho-probed sites identified after extended kinase reaction times. Of the 193 tyrosine residues, 96 were detected as phospho-probed sites following prolonged incubation of the trypsin-digested sample **(Fig. S2d)**, suggesting that these sites were phosphorylated at a slower rate due to disruption of the substrate’s higher-order structure. We also analyzed phospho-probed sites where the primary sequence surrounding the tyrosine residues was preserved; 121 sites lacked lysine or arginine in the 6 residues before and after the phosphorylation site, and their phospho-probing profiles also reflected the substrate’s higher-order structure **(Fig S2e)**.

Since the structure of proteins in the non-denatured state more closely resembles the native *in vivo* conformation than that of denatured proteins, phospho-probed sites identified in non-denatured samples are expected to reflect *in vivo* phosphorylation patterns. To assess this, we examined the overlap between phospho-probed sites identified in non-denatured samples and those reported *in vivo*. Among the tyrosine phosphorylation sites detected in non-denatured samples, 84.3 % were annotated in PhosphoSitePlus^49^, a comprehensive database of post-translational modifications. In contrast, only 65.9 % of the tyrosine phosphorylation sites uniquely identified in trypsin-digested samples were reported in the database, which is a significantly lower proportion (p < 10⁻⁵, Fisher’s exact test) **(Fig. S2f)**. These results suggest that phospho-probing of substrate proteins in the non-denatured state selectively targets phosphorylation sites that are also modified *in vivo*. In addition, tyrosine residues phosphorylated in the non-denatured state tended to be located in regions of low confidence (pLDDT) in predicted structures **(Fig. S3)**. This finding implies that phospho-probing of non-denatured substrates may provide access to regions for which current structure prediction models are less reliable.

We further quantified the phospho-probing efficiency of individual tyrosine residues in the substrate proteins and analyzed their correlation with predicted structural features. In this analysis, we defined the ratio of phospho-probed tyrosine-containing peptides in non-denatured samples to those in trypsin-digested samples (Non-denatured/Trypsin-digested ratio) as the phospho-probing efficiency for each residue. A lower ratio indicates reduced phospho-probing efficiency in the non-denatured state, while a higher ratio indicates increased efficiency. Phospho-probing efficiencies for the 247 tyrosine sites commonly identified in both conditions correlated with predicted disorder, solvent accessibility, and secondary structure of the corresponding regions **(Fig. 3d**). These results indicate that quantitative phospho-probing reflects the higher-order structural state of the substrate proteins.

We also evaluated the optimal phospho-probing reaction time for sensitively profiling differences in the higher-order structure of substrate proteins. Specifically, we investigated whether the structural dependence of phospho-probing efficiency increases or decreases with extended labeling. *In vitro* kinase reactions were performed on non-denatured and trypsin-digested samples for 15, 30, 60, and 180 minutes, and phospho-probed tyrosine residues were identified **(Fig. S4a)**. A total of 125 tyrosine phosphorylation sites were commonly detected across all time points, and the effect of incubation time on the phospho-probing efficiency of these sites was examined. We observed that the difference in phospho-probing efficiency between tyrosines located in disordered regions and those in structured regions decreased as incubation time increased **(Fig. S4b)**. This result suggests that brief kinase incubation times enhance the sensitivity of phospho-probing to differences in protein higher-order structure. Based on these findings, we adopted a 15-minute incubation for all *in vitro* kinase reactions to maximize the responsiveness of phospho-probing profiles to substrate structural states.

Overall, these results demonstrate that phospho-probing enables the profiling of protein conformational differences in complex biological samples.

### Phospho-probing enables the residue-specific profiling of protein conformational changes induced by RNA digestion in complex biological samples

Next, we further examined the applicability of phospho-probing to detect conformational changes in protein subpopulations, as the protease digestion perturbed the entire protein population in the biological sample in the previous experiment. Since SRC substrates are enriched in RNA-binding proteins **(Fig. S5)**, we anticipated that this approach would allow us to monitor protein conformational changes in response to perturbations targeting cellular RNA. Accordingly, we used phospho-probing to profile changes in the higher-order structure of proteins induced by the loss of protein–RNA interactions.

The phospho-probing reaction with SRC was conducted using non-denatured HEK293T cell lysates (RNA-preserved samples) or lysates digested with an RNase cocktail (RNase A and RNase T1) to generate RNA-digested samples **(Fig. 4a, Fig. S6a).** To improve the detectability of phosphotyrosine-containing peptides, a carrier sample enriched in tyrosine-phosphorylated peptides—generated by prolonged *in vitro* kinase reaction—was isobarically labeled and included to enhance signal detection sensitivity^39^. As a result, we successfully identified 3,126 tyrosine phosphorylation sites across 1,881 proteins. For each site, phospho-probing efficiency was calculated as the ratio of phosphorylation abundance in SRC-treated samples (SRC(+)) to that in non-treated samples (SRC(–)). Phospho-probed sites were defined as those exhibiting a statistically significant and greater than 2-fold increase in phospho-probing efficiency. Based on these criteria, 307 phospho-probed sites were identified in RNA-preserved samples and 689 in RNA-digested samples **(Fig. S6b)**. Subsequent analyses focused on 711 tyrosine residues from 587 proteins that were phospho-probed in at least one of the two conditions. Comparison of phospho-probing efficiency between RNA-preserved and RNA-digested samples revealed that 301 sites showed more than 2-fold increase in probing efficiency following RNA digestion **(Fig. 4b, Fig. S6c, S6d)**. Notably, more than 40 % of the phospho-probed tyrosine residues showed increased probing upon RNA digestion, suggesting that tyrosine residues involved in protein–RNA interactions are widely distributed across the proteome.

**Figure 4.**
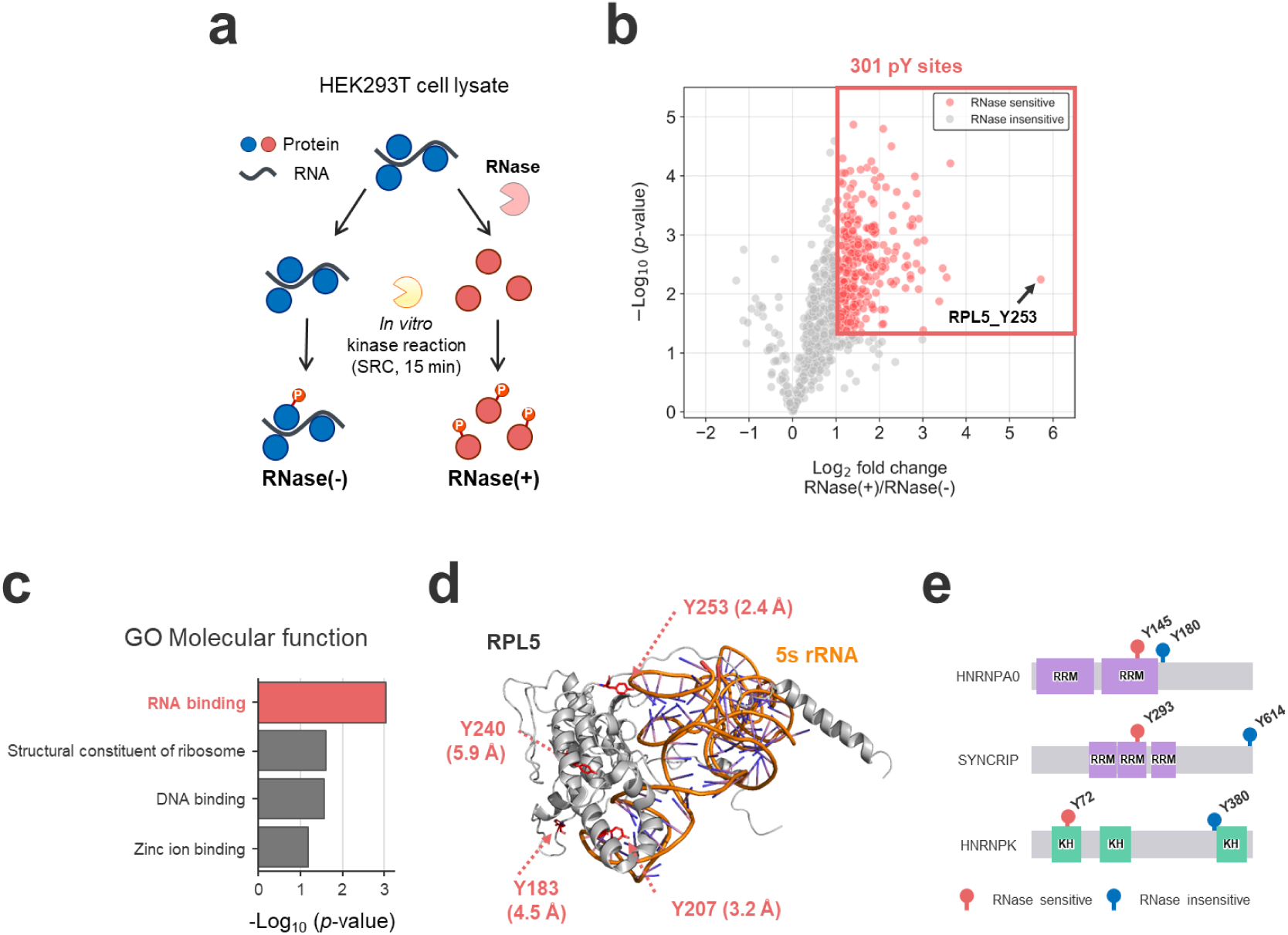
Application of phospho-probing to detect protein conformational changes induced by RNA digestion. (a) Experimental workflow for phospho-probing using RNA-preserved [RNase(–)] and RNA-digested [RNase(+)] HEK293T cell lysates. Phospho-probing reactions were performed in triplicate. (b) Volcano plot showing differences in tyrosine phospho-probing efficiency between RNA-preserved and RNA-digested samples. The x-axis represents the log₂-transformed fold change in phospho-probing efficiency following RNase treatment for 711 tyrosine phospho-probed sites. The y-axis indicates the p-value from a two-tailed Welch’s t-test. 301 sites that showed a significant increase (fold change ≥ 2; p < 0.05) in phospho-probing efficiency are highlighted in red. (c) Gene Ontology (GO) molecular function enrichment analysis of proteins containing 246 RNase-sensitive tyrosine residues. A total of 587 proteins with phospho-probed tyrosines were used as the background set. (d) Mapping of RNase-sensitive tyrosine residues onto the RPL5 and 5S rRNA complex structure (PDB: 4UG0). Tyrosine residues with increased phospho-probing efficiency in RNA-digested samples (Y183, Y207, Y240, Y253) are indicated in red. The minimum Euclidean distance (in angstroms) between each tyrosine and RNA is indicated. (e) Visualization of RNase-sensitive (red) and RNase-insensitive (blue) tyrosine residues on representative RNA-binding proteins (HNRNPA0, SYNCRIP, and HNRNPK). Schematic representations of protein domains are based on UniProt annotations. RRM: RNA recognition motif; KH: K homology domain.

We then investigated the relationship between protein–RNA interactions and tyrosine residues whose phospho-probing efficiency increased upon RNA digestion. Gene Ontology (GO) enrichment analysis using the Database for Annotation, Visualization and Integrated Discovery (DAVID)^50,51^ revealed a strong enrichment of RNA-binding proteins among the 246 RNase-sensitive proteins, indicating that phospho-probing is capable of detecting higher-order conformational changes in RNA-binding proteins **(Fig 4c)**. We further focused on 60S ribosomal protein L5 (RPL5), which contains Y253—the tyrosine residue that showed the largest increase in phospho-probing efficiency following RNA digestion among all RNA-binding proteins. In total, 4 tyrosine residues with increased phospho-probing efficiency were identified in RPL5. These sites were mapped onto the crystal structure of the protein–RNA complex, and their spatial proximity to RNA was evaluated. All 4 tyrosine residues were located within 8 Å of the RNA, indicating that they are situated close to RNA in the native complex^52,53^ **(Fig 4d)**. These findings demonstrate that phospho-probing analysis can detect sites of conformational change in proteins associated with the loss of protein-RNA interactions. We further investigated the relationship between domain localization of phospho-probed tyrosine residues and their RNase sensitivity. Among the 711 phospho-probed tyrosine residues, 125 were located within UniProt-annotated domains, of which 58 (46 %) were RNase-sensitive. Notably, 30 phospho-probed tyrosine residues were located within RNA recognition motifs (RRMs), of which 21 (70 %) were RNase-sensitive, indicating that the majority of tyrosines located within RNA-binding domains are RNase-sensitive. For example, in HNRNPA0 and SYNCRIP, tyrosines located on the RRM (HNRNPA0_Y145 and SYNCRIP_Y293) were RNase-sensitive, whereas tyrosines located outside the RRM (HNRNPA0_Y180 and SYNCRIP_Y614) did not exhibit RNase sensitivity. Similarly, in HNRNPK, the tyrosine located on the KH domain (Y72) showed RNase sensitivity, while the tyrosine outside the KH domain (Y380) did not **(Fig. 4e)**. These examples suggest that phospho-probing is capable of detecting protein conformational changes in a site-specific manner.

To gain insight into the *in vivo* function of RNase-sensitive tyrosine residues, we analyzed a phosphorylation site dataset from the literature to investigate phosphorylation within RNA-binding domains. Ramasamy *et al.* previously compiled a comprehensive dataset containing 81,404 phosphorylation sites across the human proteome through large-scale reanalysis of publicly available proteomics data^54^. This dataset includes 61 tyrosine phosphorylation sites located within RNA recognition motifs (RRMs). Of these, 11 sites were phospho-probed using SRC, and 6 exhibited increased phospho-probing efficiency following RNA digestion. For example, SNRNP70, a subunit of the small nuclear ribonucleoprotein (snRNP) complex, showed enhanced phospho-probing of Y126 upon RNA digestion **(Fig. S7)**. Structural mapping of this tyrosine residue onto the crystal structure of the snRNP complex revealed that Y126 is located at the interaction interface between SNRNP70 and RPB2, a subunit of RNA polymerase II. These findings highlight the potential of phospho-probing to detect structural alterations at biologically functional phosphorylation sites.

### Performance comparison of phospho-probing and solubility assay

Sindhuja *et al.* previously reported that digestion of RNA in cell lysates increases the solubility of proteins involved in RNA–protein interactions, based on systematic measurements of protein solubility^55^. However, proteins that are already highly soluble under RNA-preserved conditions, or those exhibiting only modest solubility changes upon RNA digestion, are likely to have their conformational changes overlooked by solubility-based methods. To evaluate the performance of phospho-probing as a tool for detecting protein conformational changes, we compared RNase-sensitive proteins identified by phospho-probing with those identified by the solubility assay, which quantifies solubility differences based on surfactant extraction. A total of 1,507 proteins were commonly present in both datasets **(Fig. 5a)**, including 480 proteins that were phospho-labeled by SRC. Among these 480 phospho-probed proteins, 166 were identified as RNase-sensitive only by phospho-probing, 31 were uniquely identified by the solubility assay, and 43 were identified by both methods **(Fig. 5b)**. The substantially greater number of RNase-sensitive proteins detected exclusively by phospho-probing suggests that this method is highly sensitive for detecting conformational changes in phospho-labeled proteins.

**Figure 5.**
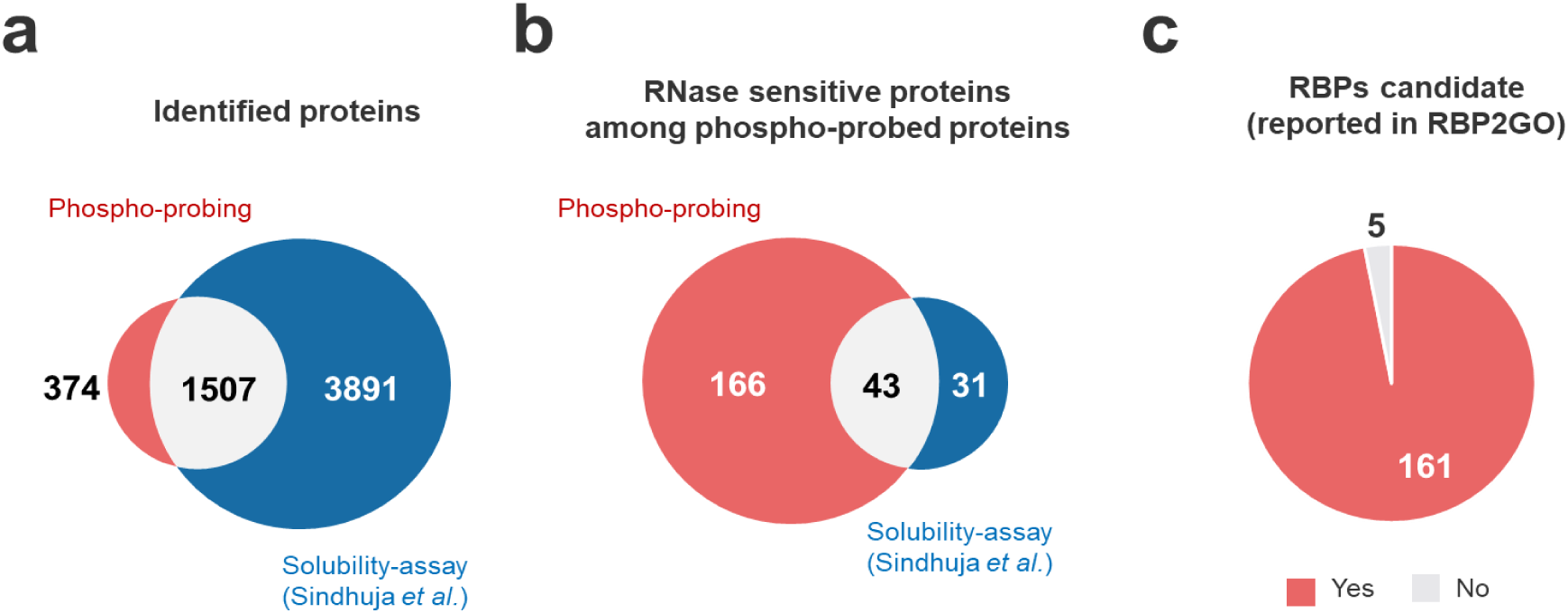
Comparison of RNase-sensitive proteins detected by phospho-probing and solubility assay. (a) Overlap of all identified proteins between phospho-probing and solubility assay datasets. (b) Overlap of RNase-sensitive proteins identified by each method among the 480 phospho-probed proteins within the 1,507 proteins commonly analyzed by both approaches. (c) Pie chart showing the proportion of the 166 RNase-sensitive proteins uniquely identified by phospho-probing that are annotated as RNA-binding proteins (RBPs) in the RBP2GO database.

Most of the 166 RNase-sensitive proteins uniquely identified by phospho-probing are annotated in RBP2GO^56^, a database of RNA-binding proteins **(Fig. 5c, Fig. S8a)**, supporting their identity as RNA-interacting proteins. In addition, solubility assay results showed that these proteins were consistently either soluble (124 proteins) or insoluble (42 proteins), regardless of RNA degradation **(Fig. S8b)**. This indicates that phospho-probing can capture RNA-dependent conformational changes that are not detectable by solubility-based methods. This discrepancy may be due to the fundamental difference in resolution between the two approaches; whereas the solubility assay reflects the changes in the physicochemical properties of the entire protein, phospho-probing enables site-specific detection of local structural alterations. Interestingly, five of the 166 RNase-sensitive proteins (NUCB2, FNBP1L, AP1AR, MIER3, and PNPO) were not annotated in RBP2GO, suggesting they may represent previously unrecognized RNA-binding proteins. For instance, MIER3 (Mesoderm Induction Early Response 1, Family Member 3) is a transcriptional repressor that forms a complex with HDAC1/HDAC2 and Snail and binds to the E-cadherin promoter to repress its expression^57^. Given this regulatory role, it would be of interest to assess the RNA-binding activity of MIER3 and other candidates identified by phospho-probing. Our findings provide a valuable resource for exploring the function and mechanisms of unknown RNA-binding proteins.

On the other hand, 31 proteins were identified as RNase-sensitive exclusively by the solubility assay. In these cases, the phospho-probed sites appeared to differ from those involved in RNA-binding activity, as fewer tyrosine residues located within RNA recognition motifs (RRMs) were phosphorylated **(Fig. S8c)**. This suggests that phospho-probing may fail to detect structural changes if the conformational shift occurs in regions of the protein that are not directly accessible for labeling. For example, although SRSF1 was successfully phospho-labeled by SRC, RNase sensitivity was detected only by the solubility assay. SRSF1 is known to form a complex with RNA via its RRM domain, but the phospho-probed tyrosine residue Y170 is located far from the RRM–RNA interaction interface **(Fig. S8d)**. This implies that while Y170 resides within a domain associated with RNA binding, the local structure surrounding this residue may not undergo significant conformational change upon RNA dissociation. Therefore, phospho-probing may overlook such changes when labeling does not occur near the site of structural alteration.

In summary, phospho-probing with kinases enabled the proteome-wide and site-specific detection of conformational changes in substrate proteins that interact directly or indirectly with RNA.

### Limitations of the study

Our results demonstrate the potential of phospho-probing as a strategy for detecting protein conformational changes through *in vitro* kinase reactions applied to purified proteins or cell-extracted proteomes. However, it remains to be determined whether phospho-probing can capture conformational changes that occur within cells. Given that previous structural proteomics approaches have successfully detected intracellular protein conformational changes^6,18^, we believe that phospho-probing has the potential to expand the capabilities of structural proteomics by enabling the detection of such changes within cells with unprecedented sensitivity.

Since phosphorylation is catalyzed by protein kinases, proteins with known phosphorylation sites are, in principle, amenable to higher-order structural analysis using the phospho-probing method. Indeed, according to PhosphoSitePlus^49^, tyrosine phosphorylation sites have been reported in 57 % (11,743 of 20,601) and 84 % (17,346 of 20,601) of human proteins. However, the proteins analyzed in this study represent only a small subset of the phosphorylated proteins documented in the human proteome. In this study, we employed only SRC kinase to evaluate the feasibility of phospho-probing for detecting protein conformational changes. In principle, any kinase could be used for *in vitro* kinase reactions, and expanding the repertoire of kinases used is expected to increase both the number of phospho-probed proteins and the analytical coverage of protein higher-order structural changes using phospho-probing.

Our phospho-probing method can analyze protein conformational changes only in proteins that contain phosphoacceptor residues recognized by kinases, along with compatible surrounding amino acid sequences. Moreover, phospho-probing reactions are not feasible under conditions where kinases are denatured or inactivated. The development of kinases with engineered substrate specificity or enhanced tolerance to denaturing conditions—through protein design technologies—could expand the applicability of this method to a broader range of structural analyses.

In the current workflow, phosphate groups introduced by phospho-probing cannot be distinguished from endogenous phosphate groups due to their identical mass. Labeling efficiency was defined based on quantification relative to the non-kinase-treated group; however, a more accurate assessment of labeling efficiency is needed. Incorporating non-natural ATP analogs, such as ¹⁸O-ATP, into the phospho-probing reaction may provide a solution by enabling clear differentiation between introduced and endogenous phosphate groups.

We used a substrate pool generated by trypsin digestion as a control to evaluate the effect of protein higher-order structure on phospho-probing. However, we also found that the loss of amino acid sequences surrounding phospho-probing sites due to trypsin digestion can influence labeling efficiency. Therefore, the choice of substrate pool for control samples remains a matter for consideration. While trypsin digests were selected for their practicality in preparing control substrates, alternative proteolytic enzymes could be used to generate peptide pools that allow for phospho-probing analysis with reduced sequence bias.

We could not distinguish whether increases in phospho-probing efficiency were due to perturbation-induced changes in protein higher-order structure or secondary conformational changes resulting from the phospho-probing process itself. Although we minimized the impact of secondary effects by employing relatively brief *in vitro* kinase reaction times, these conditions may still influence the structural state of the substrate proteins, and further optimization is required. Time-course profiling of phospho-probing efficiency could help differentiate between primary higher-order structural changes and secondary alterations induced by the labeling process.

The *in vitro* kinase reactions were performed on cell extracts in the presence of high concentrations of ATP and MgCl₂, which may have altered the native higher-order structures of the substrate proteins. As a result, the structural states observed in the extracts may not accurately reflect those *in vivo*. Performing phospho-probing reactions under physiological conditions or directly in living cells could help overcome this limitation and provide a more accurate representation of *in vivo* protein conformational states.

In this study, only peptides containing a single phospho-probed site were considered; multi-phosphorylated sites were excluded from the analysis to simplify interpretation. However, capturing higher-order conformational changes at multi-phosphorylation sites is of significant interest. Future improvements in the analytical workflow would enable the inclusion of multi-phosphorylated peptides, allowing for a more comprehensive understanding of structural dynamics at these complex regulatory sites.

## Conclusions

We have developed a novel strategy for proteome-wide analysis of protein conformational dynamics based on phospho-probing using *in vitro* kinase reactions. This approach leverages the residue-specific labeling provided by the substrate recognition ability of protein kinases, combined with the sensitive detection of low-abundance phospho-labeled peptides through phosphopeptide enrichment, to enable proteome-wide and site-specific identification of protein conformational changes. Since this phospho-probing strategy should be compatible with any kinase, the application of multiple kinases and the ability to control kinase specificity will further expand its utility for analyzing a broader range of protein conformational states.

## Resource availability

### Lead contact

Further information and requests for resources and reagents should be directed to and will be fulfilled by the lead contact, Yasushi Ishihama.

### Materials availability

This study did not generate new unique reagents.

### Data availability

- The raw MS data and analysis files have been deposited with the ProteomeXchange Consortium (http://proteomecentral.proteomexchange.org) via the jPOST partner repository (https://jpostdb.org/) with the data set identifier PXD063369.
- This paper does not report original code.
- Any additional information required to reanalyze the data reported in this paper is available from the lead contact upon request.

## Acknowledgements

We would like to thank the members of the Department of Molecular Systems BioAnalysis for the fruitful discussion. We also thank the Kyoto University Innovative Support Alliance for Life Science (iSAL) for the J-820 circular dichroism spectrometer with Prof. Kenichi Kawano and the Orbitrap Exploris 480 mass spectrometer. This work was supported by JST SPRING (JPMJSP2110 to AM), JST ACT-X (JPMJAX2324 to KO), JSPS KAKENHI (23K18185, 23H04924 to YI and 25K18607 to KO), FY 2024 Researcher Exchange Program between JSPS and ETH to KO, Nakatani Foundation to KO and the Research Foundation for Pharmaceutical Sciences to KO.

## Author contributions

Conceptual design of the project was done by KO and YI. All experiments and data analysis were carried out by AM. The first draft of the manuscript was written by AM, and the final manuscripts were prepared by AM, KO and YI.

## Declaration of generative AI and AI-assisted technologies

During the preparation of this work, the authors used GPT-4 in order to improve language and readability. After using this tool, the authors reviewed and edited the content as needed and take full responsibility for the content of the publication.

## Supplemental information

- Document S1. Figures S1-S8
- Table S1. All experimental information of phospho-probing (substrate amount, reaction time, TMT channel etc.), related to STAR methods

## STAR★Methods

### EXPERIMENTAL MODEL AND SUBJECT DETAILS

#### Apomyoglobin preparation

Apomyoglobin was prepared by removing heme from holomyoglobin (Sigma) as previously described^58^. The resulting apomyoglobin solution was exchanged to reaction buffer [40 mM Tris-HCl (pH 7.5), 20 mM MgCl₂] using a 3-kDa ultrafiltration device (Amicon Ultra, Millipore). The final protein concentration was adjusted to 1 mg/mL. The concentration of apomyoglobin was estimated based on the dry weight of the lyophilized holomyoglobin.

#### Cell culture

HEK293T cells (RIKEN BRC cell bank) were cultured in Dulbecco’s modified Eagle’s medium with 10 % fetal bovine serum in 10 cm or 15 cm diameter dishes at 37 °C under 5 % CO_2_ in air until 80 % confluency was reached. For harvesting, cells were washed twice with ice-cold phosphate-buffered saline (PBS), detached using a cell scraper, and pelleted by centrifugation.

## METHOD DETAILS

### Thermal denaturation of myoglobin

50 μL aliquots of apomyoglobin solution were transferred to a 0.2 mL polymerase chain reaction (PCR) tube and heated in parallel at 37 °C, 42 °C, 47 °C, 52 °C, 57 °C, 62 °C, 67 °C, and 72 °C for 15 minutes using the VeritiPro Thermal Cycler (Thermo Fisher Scientific). After heating, the samples were placed on ice.

### Circular dichroism (CD) spectroscopy

CD spectra were recorded using a J-820 spectropolarimeter (Jasco). The heated protein solution was diluted 5-fold with reaction buffer, and far-UV CD spectra were acquired at 37 °C in the range of 200 to 250 nm. A quartz cell with a 1 mm path length was used for measurements.

### Protein extraction from cells and RNA digestion treatment

HEK293T cell pellets were resuspended in ice-cold lysis buffer [40 mM Tris-HCl (pH 7.5), protease inhibitor cocktail (Sigma), phosphatase inhibitor cocktail 2 and 3 (Sigma), 1 U/mL RNasin® Plus Ribonuclease Inhibitor (Promega)]. The suspension was passed 10 times through a 22G needle and 3 times through a 26G needle, then centrifuged at 20000 × g for 20 minutes at 4 °C using a benchtop centrifuge (Eppendorf). The protein concentration in the resulting supernatant was measured using the Pierce BCA protein assay (Thermo Fisher Scientific) and diluted to 1 mg/mL with lysis buffer. The extracted protein solution was divided into two aliquots. One aliquot was used as the RNA-preserved lysate, and the other one was used to prepare RNA-digested lysate. RNA digestion was performed based on the method described by Sindhuja et al^55^. For the digestion, 2 μL of RNase cocktail (Thermo Fisher Scientific) was added per 100 μL of protein solution, followed by incubation at 4 °C for 30 minutes.

### *In vitro* kinase reaction of substrate proteins

The *in vitro* kinase reaction was performed using recombinant tyrosine kinase SRC (human, Carna Biosciences) at an enzyme-to-substrate (E:S) ratio of 1/1000 (w/w) in the presence of 1 mM ATP and 20 mM MgCl₂ at 37 °C for 15 minutes. For evaluation of the SRC kinase activity toward myoglobin, the reaction was extended to 3 hours. For evaluation of the optimal phospho-probing reaction time, the reaction was performed for 15, 30, 60 and 180 minutes. The kinase reaction was quenched by the addition of sodium deoxycholate (final concentration: 12 mM) and sodium N-lauroylsarcosine (final concentration: 12 mM), followed by heat treatment at 95 °C for 5 minutes. The number of replicates for each *in vitro* kinase reaction is listed in **Supplementary Table S1.**

### Protein digestion and desalting

Protein digestion and desalting were performed based on the sample preparation method using phase transfer surfactant (PTS)^59^. Proteins were reduced with 10 mM dithiothreitol (DTT) for 30 minutes and alkylated with 50 mM iodoacetamide (IAA) for 30 minutes in the dark. The sample solution was diluted fivefold with 50 mM ammonium bicarbonate, and Lys-C (Wako) and trypsin (Promega) were added at an E:S ratio of 1:100 (w/w) each, followed by overnight digestion at 37 °C. After digestion, an equal volume of ethyl acetate was added and acidified with 0.5 % trifluoroacetic acid (TFA). After shaking for 2 minutes, the sample was centrifuged at 15,800 × g for 2 minutes. The aqueous and organic phases were completely separated, and the aqueous phase was dried using a SpeedVac (Thermo Fisher Scientific), reconstituted in 0.1 % TFA and 5 % acetonitrile (ACN), and desalted using SDB-XC StageTips^60–62^.

### *In vitro* kinase reaction of tryptic digests

Desalted peptides were dried and then reconstituted in 40 mM Tris-HCl (pH 7.5) at 1 mg/mL. Note that the peptide amount corresponds to the protein amount measured by the BCA assay prior to digestion. The *in vitro* kinase reaction was performed using recombinant tyrosine kinase SRC at an E:S ratio of 1/1000 (w/w) in the presence of 1 mM ATP and 20 mM MgCl₂ at 37 °C for 15 minutes. For characterization of the phospho-probed sites uniquely identified in non-denatured samples, the reaction was performed for 60 minutes. For evaluation of the optimal phospho-probing reaction time, the reaction was performed for 15, 30, 60 and 180 minutes. For preparation of the carrier sample used for isobaric labeling signal amplification, the *in vitro* kinase reaction was performed at an E:S ratio of 1/500 (w/w) for 3 hours. The kinase reaction was quenched by acidifying the solution to a final concentration of 0.5 % TFA, and the peptides were desalted using SDB-XC StageTips. The number of replicates for each *in vitro* kinase reaction is listed in **Supplementary Table S1.**

### Phosphopeptide enrichment

Phosphopeptide enrichment was performed using hydroxy acid-modified metal oxide chromatography (HAMMOC)^31^ . Briefly, TiO₂ beads were packed into a StageTip equipped with an Empore C8 disc (GL Sciences) and coated with lactic acid. The digested peptides were then loaded onto the StageTip, allowing phosphorylated peptides to coordinate with the TiO₂ beads. Non-phosphorylated peptides that interacted weakly with TiO₂ were removed by washing. Phosphopeptides were subsequently eluted using 0.5 % piperidine, acidified with phosphoric acid, and desalted using an SDB-XC StageTip. The desalted peptides were dried using a SpeedVac and reconstituted in the LC/MS/MS solvent containing 0.5 % TFA and 4 % ACN. For stable isotope labeling experiments, heptafluorobutyric acid (HFBA) was added instead of phosphoric acid before introducing the sample into the StageTip-based microreactor. The information on the peptide amount in each phosphopeptide enrichment for all experiments is shown in **Supplementary Table S1**.

### TMT labeling

Isobaric labeling of phosphopeptides was performed using TMTpro (Thermo Fisher Scientific) by a solid-phase reaction method^63^. SDB-XC StageTips packed with 0.5 mg of InertSep® RP-C18 beads (GL Sciences) were activated with 0.1 % TFA and 80 % ACN and equilibrated with 0.1 % TFA and 5 % ACN. Phosphopeptides enriched from 50 μg of protein were loaded onto the StageTips and washed with 50 mM phosphate buffer (pH 6.5). To avoid peptide elution, TMT reagents were dissolved in ACN and then diluted with phosphate buffer to reduce the ACN concentration to less than 5 %. A volume of 10 μL of this TMT solution (10 μg/10 μL) was applied to each tip. The particles were kept moist, and samples were incubated at room temperature for 1 hour. The StageTips were then washed with 0.1 % TFA and 5 % ACN, and peptides were eluted with 0. 1 % TFA and 80 % ACN. The information on the peptide amount in each TMT channel is shown in **Supplementary Table S1**.

### High-pH reverse-phase chromatography-based fractionation

High-pH reverse-phase chromatography-based fractionation of TMT-labeled phosphopeptides was performed using StageTips as previously described^64^ with minor modifications. TMT-labeled phosphopeptides were pooled and dried in a SpeedVac, then reconstituted in 50 μL of 0.1 % ammonia. SDB-XC StageTips were activated with methanol, 0.1 % ammonia, and 80 % ACN, followed by equilibration with 0.1 % ammonia. The phosphopeptides were then loaded onto the StageTip, and the flow-through fraction was collected. Peptides were sequentially eluted with 0.1 % ammonia containing 5 %, 7.5 %, 10 %, 12.5 %, 15 %, 20 %, and 80 % ACN. The flow-through and the first fraction (5 %) were pooled, as were the last two fractions (20 % and 80 %). The recovered peptides were dried using a SpeedVac and reconstituted in solvent consisting of 0.5 % TFA and 4 % ACN for nanoLC/MS/MS analysis.

### NanoLC/MS/MS analysis

The peptide digests of myoglobin were analyzed using a nanoLC/MS/MS system consisting of a PAL HTC-xt autosampler (CTC-Analytics), an UltiMate 3000 RSLCnano pump (Thermo Fisher Scientific), and a Q Exactive mass spectrometer (Thermo Fisher Scientific). The mass spectrometric analyses were carried out in the data-dependent acquisition (DDA) mode. The LC mobile phases consisted of solvent A (0.5 % acetic acid) and solvent B (0.5 % acetic acid and 80 % ACN). Peptides were separated using an in-house-packed needle column^65^ (150 mm, 100 μm i.d.) containing Reprosil-Pur 120 C18-AQ 3.0 μm (Dr. Maisch). The flow rate was set to 500 nL/min. The gradient conditions for mobile phase B were as follows: 5–10 % B in 0.5 min, 10–40 % B in 20 min, 40–99 % B in 1 min, followed by 99 % B for 3.5 min. The electrospray voltage was set to 2.4 kV in the positive mode. The mass range of the survey scan was from 350 to 1500 m/z with a resolution of 70,000, an automatic gain control (AGC) target of 3 × 10⁶, and a maximum injection time (MIT) of 100 ms. The first mass of the MS/MS scan was set automatically with a resolution of 17,500, an AGC target of 1 × 10⁵, and a MIT of 100 ms. The fragmentation was performed by higher energy collisional dissociation with a normalized collision energy of 27 %. The dynamic exclusion time was set to 30 s.

For HEK293T proteome samples, peptide digests were analyzed using a nanoLC/MS/MS system consisting of a WPS-3000PL RS autosampler (Thermo Fisher Scientific), an UltiMate 3000 RSLCnano pump, a FAIMS Pro interface (Thermo Fisher Scientific), and an Orbitrap Exploris 480 mass spectrometer (Thermo Fisher Scientific). The mass spectrometric analyses were carried out in the DDA mode. The LC mobile phases consisted of solvent A (0.5 % acetic acid) and solvent B (0.5 % acetic acid and 80 % ACN). Peptides were separated using an in-house-packed needle column (250 mm, 100 μm i.d.) containing Reprosil-Pur 120 C18-AQ 1.9 μm resin (Dr. Maisch) at 50 °C in a PRSO-V2 column oven (Sonation GmbH). The flow rate was set to 400 nL/min. The gradient conditions were as follows: 5 % B in 5 min, 5–19 % B in 55.3 min, 19–29 % B in 21 min, 29–40 % B in 8.7 min, 40–99 % B in 0.1 min, followed by 99 % B for 4.9 min. The electrospray voltage was set to 2.4 kV in the positive mode. The FAIMS mode was set to standard resolution, and the total carrier gas flow was 4.0 L/min. The CV was set to −40 and −60, and the cycle time of each CV experiment was set to 1 s. The mass range of the survey scan was from 375 to 1500 *m/z* with a resolution of 60,000, 300 % normalized AGC target, and auto MIT. The first mass of the MS/MS scan was set to 120 *m/z* with a resolution of 15,000, standard AGC target, and auto MIT. The fragmentation was performed by higher energy collisional dissociation with a normalized collision energy of 30 %. The dynamic exclusion time was set to 20 s.

For TMT-labeled proteome samples, analysis conditions were essentially the same as above, with the following changes: the gradient conditions were as follows: 5 % B in 5 min, 5–12.5 % B in 5 min, 12.5–19 % B in 21 min, 19–29 % B in 31 min, 29–40 % B in 13.7 min, 40– 99 % B in 0.1 min, followed by 99 % B for 4.9 min. The cycle time of each CV experiment was set to 1.5 s. The mass range of the survey scan was from 375 to 1575 *m/z* with a resolution of 60,000, 300 % normalized AGC target, and a MIT of 50 ms. The first mass of the MS/MS scan was set to 110 *m/z* with a resolution of 45,000, 200 % normalized AGC target, and a MIT of 120 ms. The fragmentation was performed by higher energy collisional dissociation with a normalized collision energy of 32 %. The dynamic exclusion time was set to 30 s.

## QUANTIFICATION AND STATISTICAL ANALYSIS

### MS data processing

For myoglobin datasets, peptides were identified by automated database searches with Mascot ver. 2.7.0 (Matrix Science) against the horse myoglobin FASTA database (ver. 2020-05), including the MSFragger contaminants fasta database. The peptide mass tolerance was set to 5 ppm, and the fragment ion mass tolerance was set to 20 ppm. Trypsin was specified as the digestion enzyme, allowing up to two missed cleavage sites. Carbamidomethylation (C) was set as a fixed modification, while oxidation (M) and phosphorylation (STY) were set as variable modifications. Peptides were accepted if the Mascot score was over the 95 % confidence limit (p < 0.05) based on the identity score of each peptide. For HEK293T proteome datasets, the raw data files were analyzed and processed using MaxQuant (ver. 1.6.17.0)^66^, and peptides and proteins were identified by automated database searches with Andromeda^67^ against the UniProt human fasta file (ver. 2022-03, 20,601 human protein entries). Raw data files collected from FAIMS experiments were split into a set of MaxQuant compliant MzXML files using FAIMS MzXML Generator (https://github.com/coongroup/FAIMS-MzXML-Generator). The precursor mass tolerance was set to 4.5 ppm, and the fragment ion mass tolerance was set to 20 ppm. Trypsin and LysC were specified as the digestion enzymes, allowing up to two missed cleavage sites. For label-free quantification experiments, carbamidomethylation (C) was set as a fixed modification, and oxidation (M) and phosphorylation (STY) were set as variable modifications. For TMT quantification experiments, TMTpro (peptide N-term and K) and carbamidomethylation (C) were set as fixed modifications. The false discovery rate (FDR) cut-off of 1 % was applied for both peptide-spectrum match (PSM) and protein identifications. The minimum Andromeda score cut-off of 40 was applied for modified peptide identifications. For label-free quantification, the match-between-run function was used with default settings (0.7 min match window and 20 min alignment time).

### Quantification of phospho-probing efficiency of myoglobin

Identified myoglobin peptides were quantified by processing MS1 extracted ion chromatograms using Skyline (ver. 21.2.0)^68,69^. The phospho-probing efficiency of myoglobin was calculated based on the average quantification values of phosphorylated peptides from heat-denatured myoglobin.

### Quantification of phospho-probing efficiency in cell lysate

Phospho-probing efficiency in cell lysates was calculated using the *evidence.txt* output file from MaxQuant. The *Phospho(STY)Sites.txt* file was not used because quantitative values for individual phosphorylation sites may be derived from multiple distinct peptides. Similarly, entries in the *modificationSpecificPeptides.txt* file may contain phosphorylation sites at different positions within the same peptide sequence.

The calculation of phospho-probing efficiency was performed using Python (version 3.12.3) based on the *evidence.txt* file exported from MaxQuant. The data were filtered to exclude entries corresponding to the reverse decoy database, potential contaminants, and peptides with phosphorylation site localization probabilities below 0.75. Only monophosphorylated peptides were used for further analysis. In TMT-based quantification, an additional filter was applied, as previously described^39^, to minimize the ion sampling effect^70,71^; the ratio of the total TMT signal in the carrier channel to the total TMT signal in the sample channel was less than 100. To correct for differences in MS injection amounts across samples, the quantification values of tyrosine-phosphorylated peptides were normalized to the median intensity of identified serine-phosphorylated peptides. For label-free quantification experiments, the quantitative value of the phosphorylation site was based on the peptide species with the largest sum of the intensities of the phosphorylated peptides at the same position in the same peptide sequence. For TMT experiments, the quantitative value of phosphorylation sites was based on the peptide species with the lowest posterior error probability (PEP) among those with valid reporter ion signals across all sample channels. Tyrosine phospho-probing efficiency was calculated as the ratio of the mean quantification value of tyrosine phosphorylation sites in the kinase-treated sample to that in the non-treated sample.

### Volcano plot

Volcano plots were generated based on the log₂ fold change (FC) in the average quantitative values of tyrosine phosphorylation sites (x-axis) and the –log₁₀ p-values from Welch’s two-tailed t-test (y-axis). For differential analysis between kinase-treated and non-treated samples, the quantification values of tyrosine phosphorylation sites in kinase-treated and non-treated samples were used for statistical testing. Tyrosine sites with log₂ FC > 1 and p < 0.05 were considered significantly phospho-probed *in vitro*. For differential analysis of phospho-probing efficiency between RNA-digested and RNA-preserved samples, the SRC(+)/SRC(–) ratios in each sample were used for statistical testing. Tyrosine phosphorylation sites with log₂ FC > 1 and p < 0.05 were considered to exhibit a significant increase in phospho-probing efficiency upon RNA digestion.

### Annotation of structural information on phospho-probed sites

Annotation of the structural information of the identified phospho-probed sites was performed using the StructureMap python package^47^. Of the proteins in the UniProt human protein FASTA file (2022-03 release, 20,601 proteins), 20,220 predicted structures were downloaded from the AlphaFold database. Those that could not be downloaded were proteins for which AlphaFold did not provide predicted structures, such as proteins with sequence lengths longer than 2,700 and proteins containing non-canonical amino acids. Using the downloaded predicted structures, we obtained the predicted structure confidence level (pLDDT score) of the tyrosine residue and information about the protein secondary structure where the tyrosine residue is located, then the prediction-aware partially sphere exposure (pPSE) was calculated and the intrinsically disordered region (IDR) was predicted. The obtained structural information was matched to the identified phosphorylated receptor residues based on the Uniprot accession ID and residue number.

### Integration of other databases

Known phosphorylation site information was annotated using PhosphoSitePlus (https://www.phosphosite.org/). Among the phosphorylation sites listed in the “Phosphorylation site dataset” (downloaded in 2023-08), sites reported in human proteins were defined as “Reported”. Gene Ontology (GO) annotation and GO enrichment analysis of the identified proteins were performed using DAVID (https://davidbioinformatics.nih.gov/). Protein domain information corresponding to the identified phospho-probed sites was annotated using the “Add sequence features” function in Perseus (ver. 2.0.1.1)^72^. Analysis of RNA-binding domains known to be phosphorylated *in vivo* was conducted using a high-confidence, large-scale phosphorylation site dataset filtered by Ramasamy *et al.* (pr2c00164_si_001.xlsx)^54^. Protein domain information for phosphorylation sites in this dataset was also annotated using Perseus (ver. 2.0.1.1), and the dataset was matched to the identified phospho-probed sites. A comprehensive dataset of RNA-binding proteins was obtained from RBP2GO (https://rbp2go.dkfz.de/). The human RNA-binding protein dataset (downloaded in 2024-05; 6,100 proteins) from RBP2GO was matched to the identified proteins.

### Comparison with solubility-assay data

For comparison with the solubility-assay data reported by Sindhuja *et al.*, we downloaded the supplementary data containing protein solubility profiles in RNA-preserved and RNA-digested lysates from Sindhuja *et al*. 2022 (41589_2022_1062_MOESM2_ESM.xlsx; 41589_2022_1062_MOESM17_ESM.xlsx). The first identifier of “gene_name” used in the supplemental data was mapped to Uniprot accession IDs and matched to the proteins identified in this study. Based on the criteria described in the original publication^55^, solubility-assay target proteins were classified as “RNase-sensitive,” “RNase-insensitive,” or “Predominantly soluble”.

### Mapping of phospho-probed sites on protein 3D structures

Phospho-probed sites were mapped onto representative 3D protein structures obtained from either the PDB database (https://www.rcsb.org/) or the AlphaFold database (https://alphafold.ebi.ac.uk/). Protein structures were visualized using PyMOL (https://pymol.org/). The structures of horse myoglobin, RPL5, SNRNP70, and SRSF1 were retrieved from the PDB. For RPL5, the minimum Euclidean distance between tyrosine residues that exhibited increased phospho-probing efficiency upon RNA digestion and nearby RNA molecules was calculated using the “distance” function in PyMOL. Predicted structures for EIF3D and HNRNPL were obtained from the AlphaFold database.

### Statistical analysis

Quantitative data are expressed in bar and line graphs as mean ± SD (error bar). Two-tailed Welch’s t-test, the Fligner–Killeen test and two-sided Fisher’s exact test were performed using Scipy^73^ (version 1.13.1). Mann-Whitney U test and statistical annotations on box plots were performed using statannotations^74^ (version 0.6.0).

---

**Figure S1.**
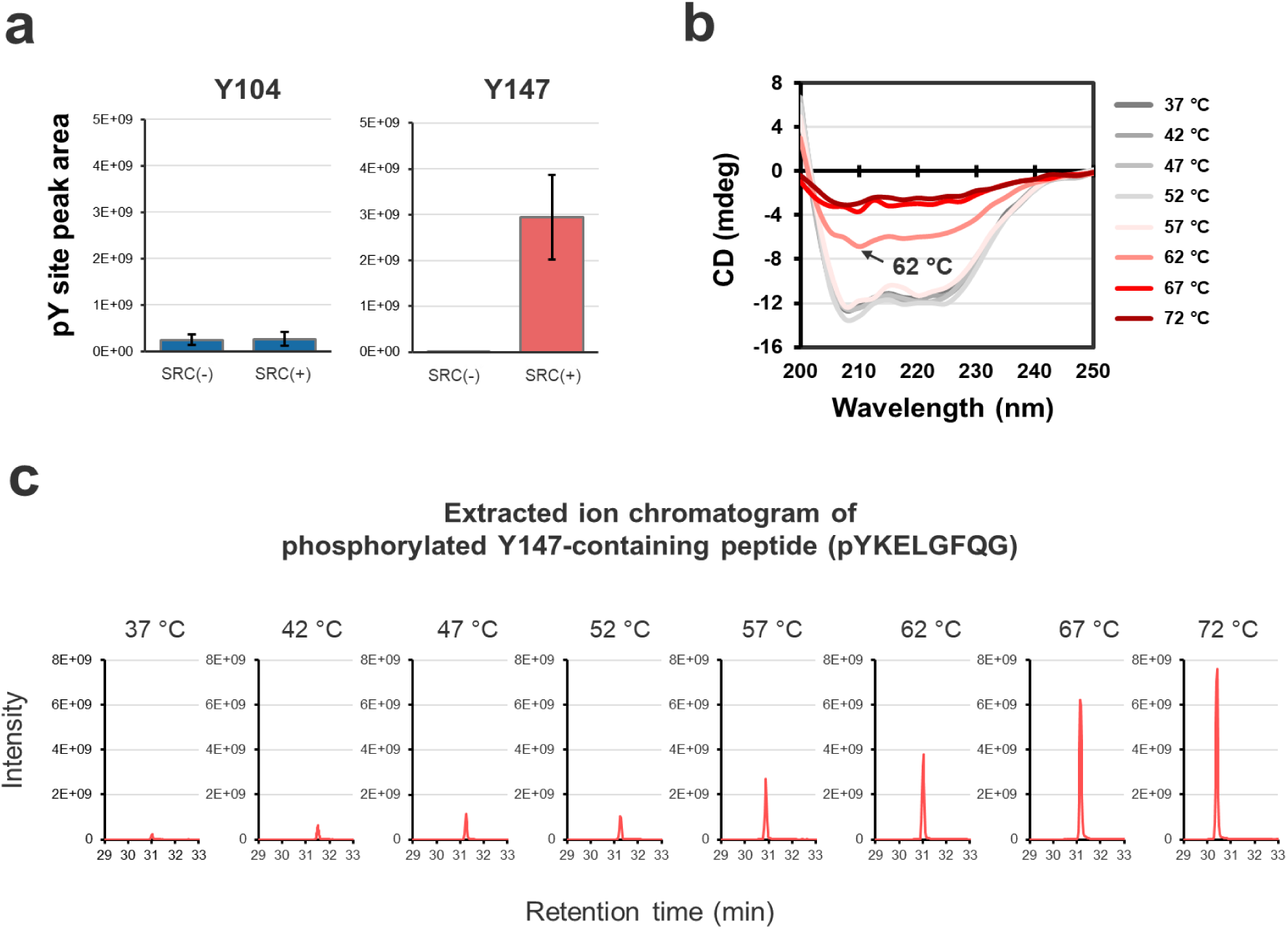
Phospho-probing using SRC kinase and secondary structure analysis of heat-denatured apomyoglobin. (a) Phospho-probing efficiency of SRC kinase on non-denatured apomyoglobin. Peak areas of tyrosine phospho-probed sites (pY sites) corresponding to the two tyrosine residues in apomyoglobin are shown for both the SRC-untreated [SRC (–)] and SRC-treated [SRC (+)] groups. Values represent the average of triplicate experiments. Error bars indicate standard deviation. (b) CD spectrum of heat-treated apomyoglobin. Arrows indicate the spectrum corresponding to apomyoglobin heated at 62 °C. (c) Representive extracted ion chromatogram (m/z = 511.2232) of the peptide containing phospho-probed Y147 of apomyoglobin (pYKELGFQG).

**Figure S2.**
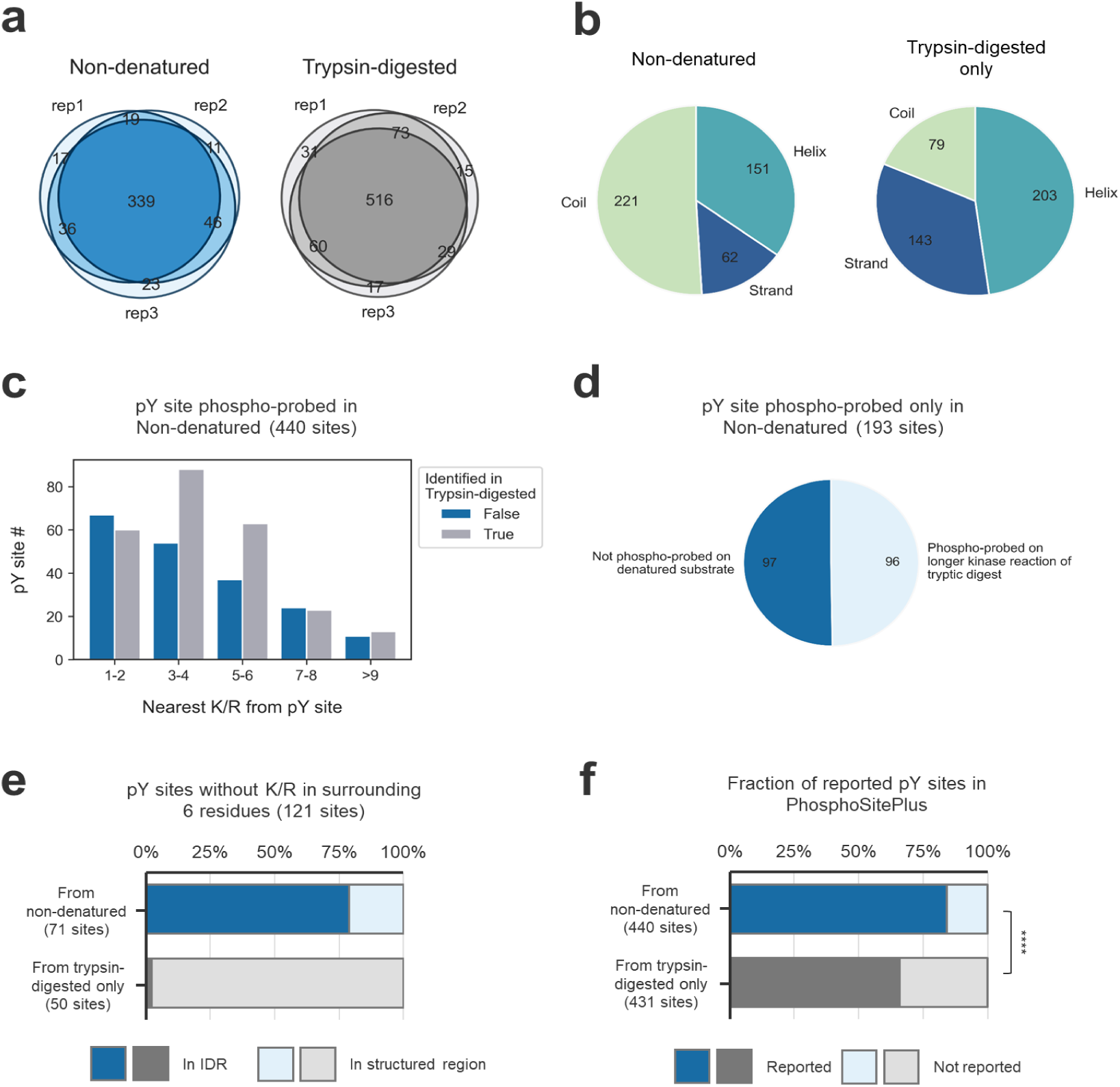
Properties of phospho-probed sites of cell lysates in the non-denatured state. (a) Overlap of tyrosine phospho-probed sites identified in non-denatured and trypsin-digested samples. Data from three technical replicates are shown. (b) Pie chart showing the distribution of predicted secondary structures, based on predicted structures in the AlphaFold database, in which tyrosine phospho-probed sites were located. (c) Distribution of lysine (K) or arginine (R) residues in the vicinity of the 440 tyrosine phospho-probed sites identified in non-denatured samples. (d) Pie chart showing the proportion of the 193 phospho-probed sites uniquely identified in non-denatured samples that were also detected by extended phospho-probing (60 min) of trypsin-digested samples. (e) Proportion of sites annotated as being located in intrinsically disordered regions (IDRs) by StructureMap, among the 121 phospho-probed sites lacking lysine or arginine in the 6 residues before and after the phosphorylation site. Tyrosine residues from proteins not available in the AlphaFold Protein Structure Database were excluded from the analysis. (f) Proportion of identified tyrosine phospho-probed sites annotated as tyrosine phosphorylation sites in the PhosphoSitePlus database. Asterisks indicate the results of Fisher’s exact test (****: p<0.0001).

**Figure S3.**
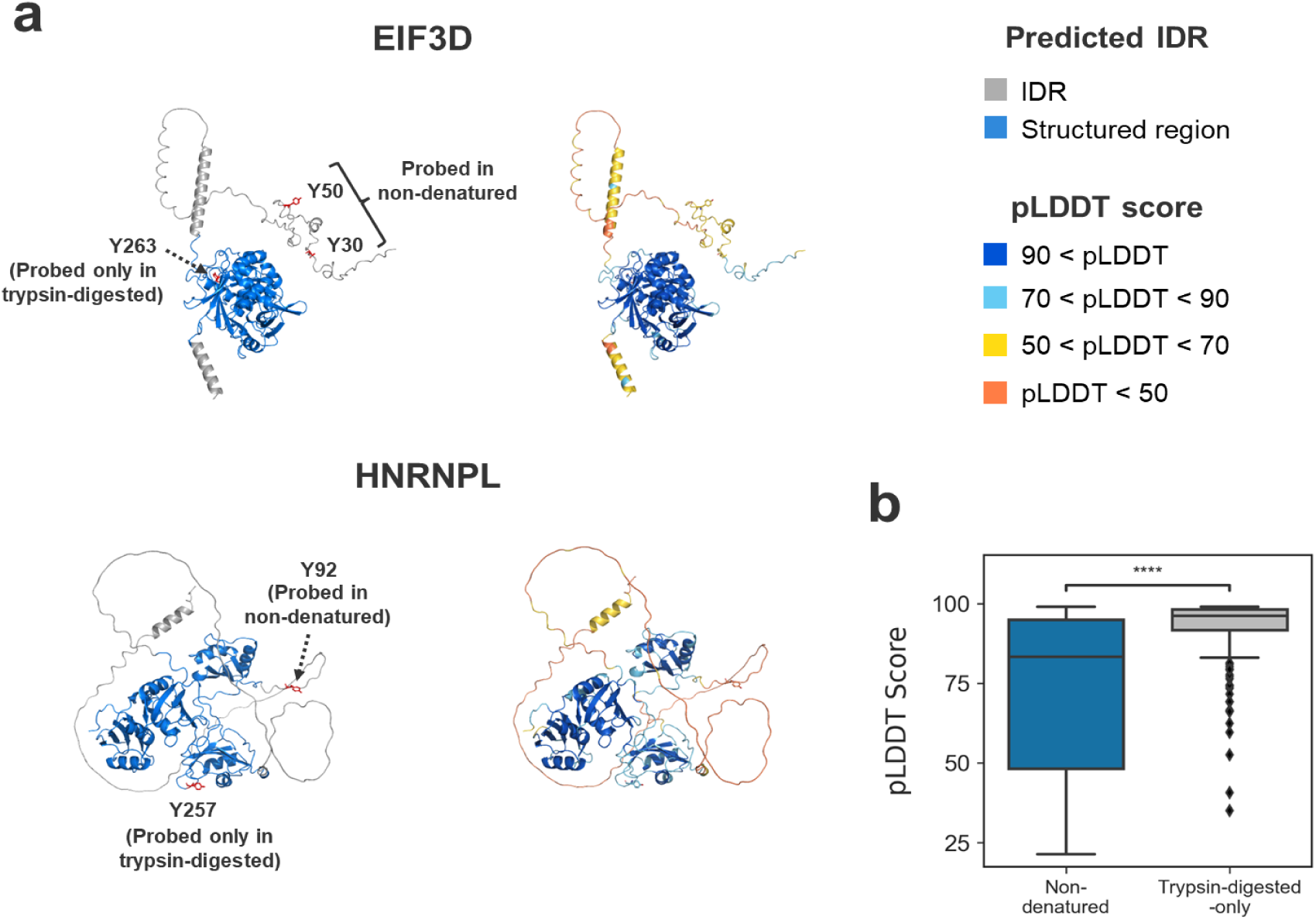
Mapping of phospho-probed sites to predicted protein structures and associated confidence scores. (a) Predicted protein structures of EIF3D (top) and HNRNPL (bottom), obtained from the AlphaFold database. Identified tyrosine phospho-probed sites are indicated in red. Protein structures are colored according to regions predicted as intrinsically disordered (IDRs) by StructureMap and by the predicted local distance difference test (pLDDT) score. In HNRNPL, Y92, which was phosphorylated in the non-denatured state, was located within an IDR, whereas Y257, which was phosphorylated in the trypsin-digested state, was located within a structured region. (b) Box plots showing the distribution of pLDDT scores for tyrosine phospho-probed sites. Sites identified in non-denatured samples had significantly lower pLDDT scores than those identified only in trypsin-digested samples. The box itself spans the interquartile range. The whiskers represent 5 % and 95 % quantiles. The thick horizontal line in each box indicates the median. Asterisks indicate results of the Mann– Whitney U test (***p < 0.0001).

**Figure S4.**
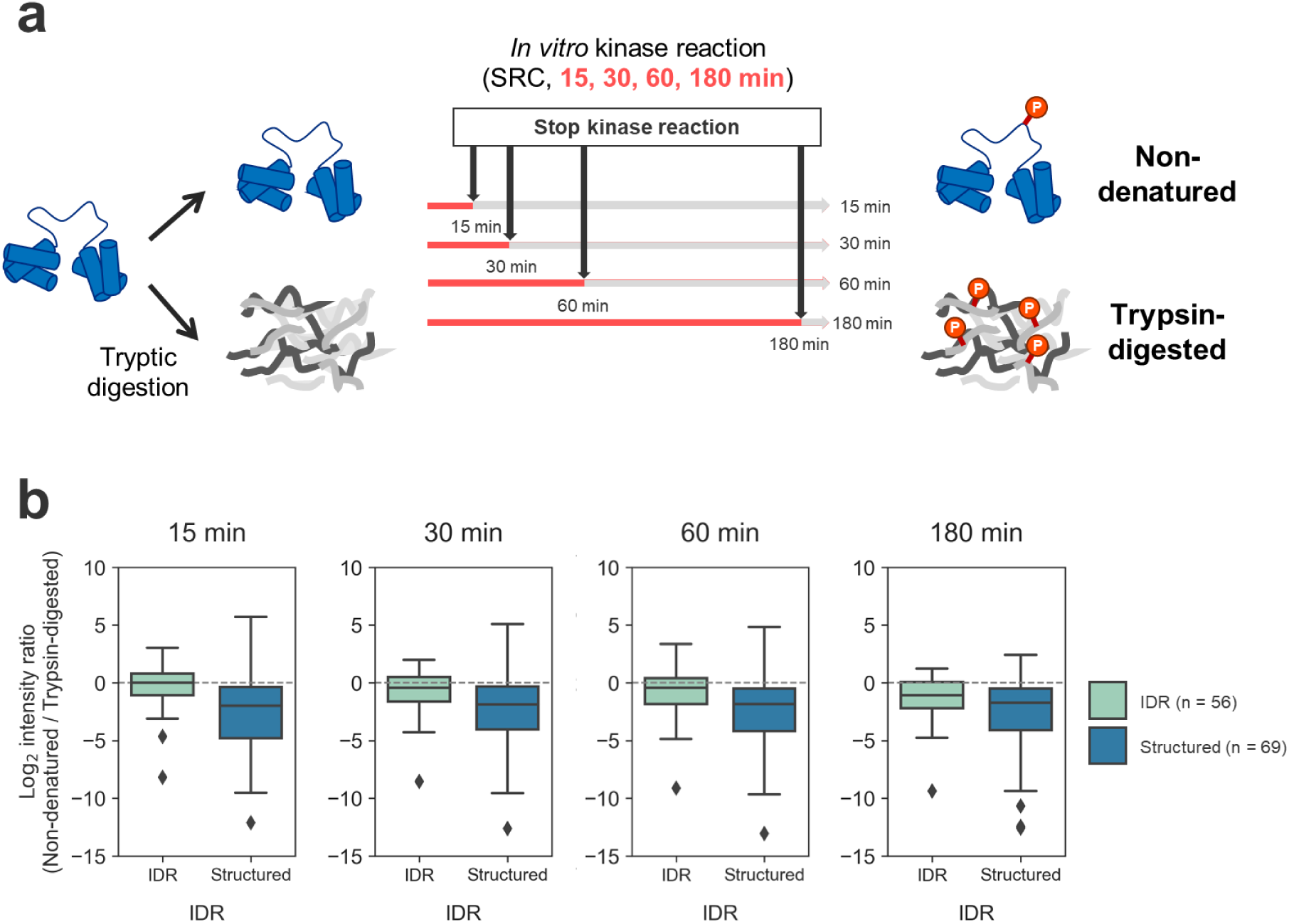
Effect of phospho-probing reaction time on the correlation between phospho-probing profiles and substrate higher-order structure. (a) Experimental workflow. Non-denatured and trypsin-digested samples were subjected to phospho-probing by *in vitro* kinase reaction. The kinase reaction was performed using recombinant tyrosine kinase SRC at an E:S ratio of 1/1000 (w/w) in the presence of 1 mM ATP and 20 mM MgCl₂ at 37 °C for 15, 30, 60 and 180 minutes. (b) Box plots showing the phospho-probing efficiency of tyrosine residues located in different predicted structural regions (IDR: n = 56, Structured: n = 69), comparing non-denatured samples to trypsin-digested samples. The box itself spans the interquartile range. The whiskers represent 5 % and 95 % quantiles. The thick horizontal line in each box indicates the median.

**Figure S5.**
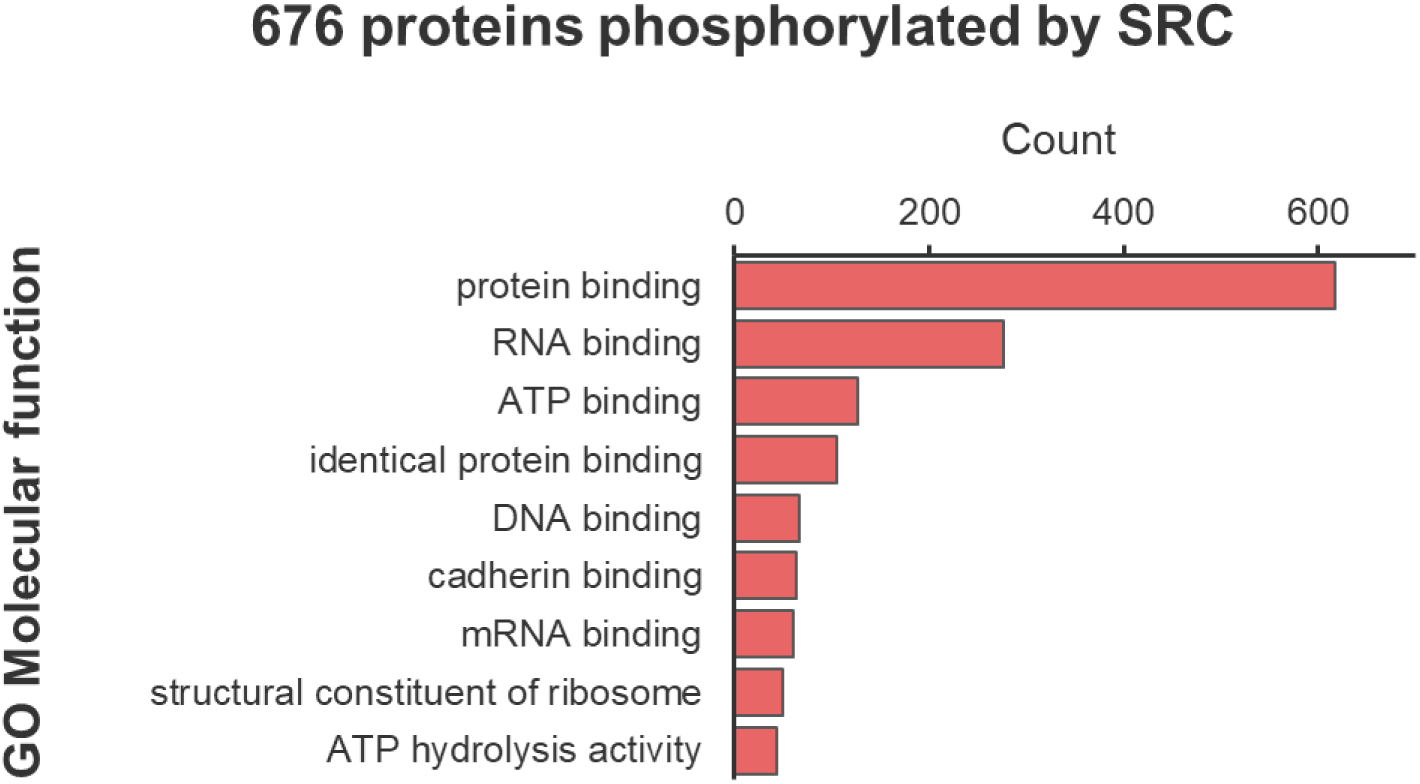
Molecular functions of proteins phospho-probed with SRC. Gene Ontology (GO) molecular functions were annotated for 676 proteins phosphorylated in a 15-minute *in vitro* kinase reaction using SRC. GO annotations were assigned using DAVID^50,51^.

**Figure S6.**
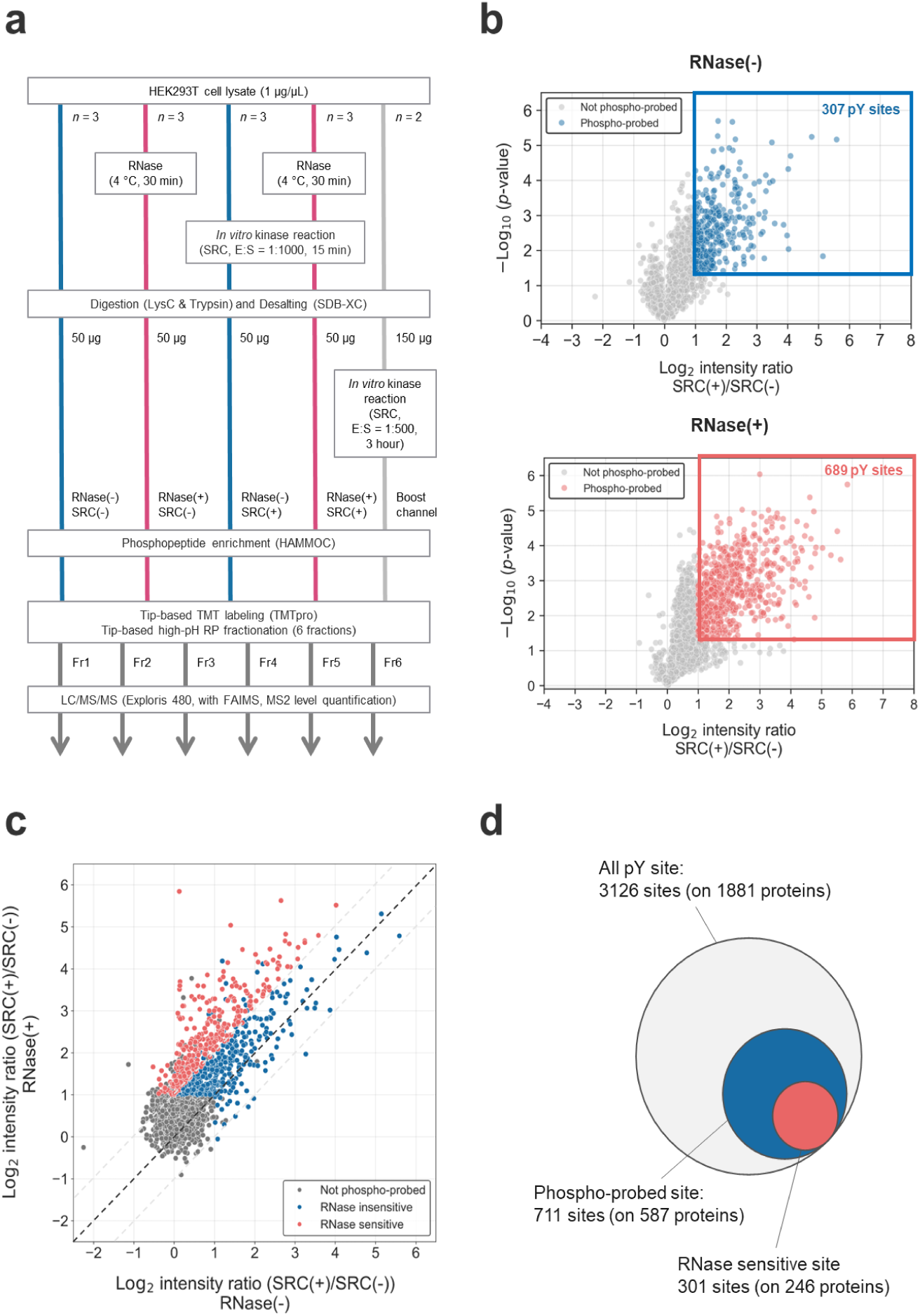
Analysis of protein conformational change due to RNA digestion by phospho-probing. (a) Detailed workflow for detecting protein conformational changes induced by RNA digestion using phospho-probing. (b) Volcano plot showing phospho-probing efficiency (SRC(+)/SRC(–) ratio) of tyrosine residues in RNA-preserved [RNase(–)] and RNA-digested [RNase(+)] samples. Tyrosine phosphorylation sites with significantly higher abundance (p < 0.05) and at least a 2-fold increase in SRC-treated samples compared to SRC-untreated controls were classified as phospho-probed sites (307 sites in RNase(–) and 689 sites in RNase(+)). (c) Scatter plot comparing phospho-probing efficiency (SRC(+)/SRC(–) ratio; log₂ scale) between RNA-preserved (x-axis) and RNA-digested (y-axis) samples. Tyrosine residues phospho-probed in at least one condition are shown in blue; those with a significant (p < 0.05, two-tailed Welch’s t-test) and more than 2-fold increase in phospho-probing efficiency upon RNA digestion are highlighted in red. (d) Summary of identified phospho-probing tyrosine sites. In total, 3,126 tyrosine phosphorylation (pY) sites were identified across 1,881 proteins. Of these, 711 sites on 587 proteins were phospho-probed by SRC. Among them, 301 sites on 246 proteins showed a significant increase in phospho-probing efficiency following RNA digestion.

**Figure S7.**
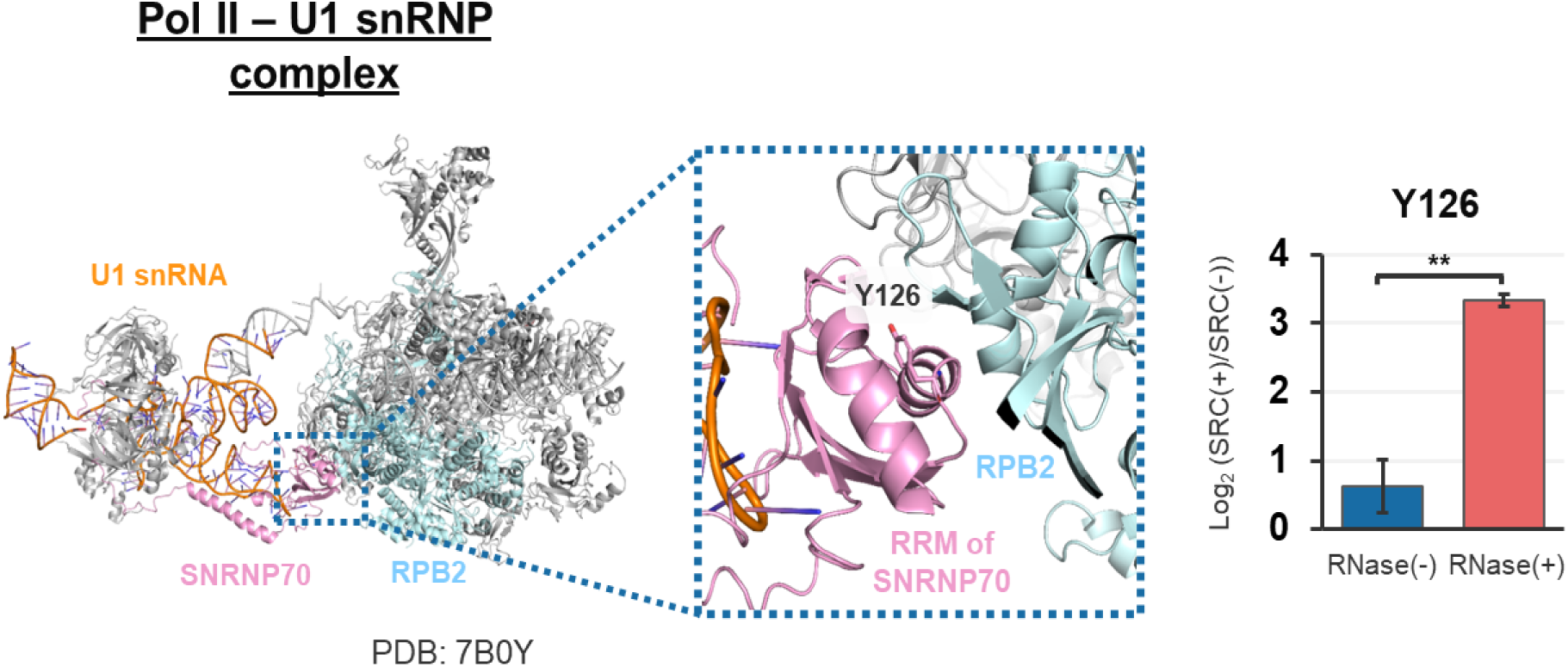
Mapping of RNase-sensitive tyrosine in the transcribing RNA polymerase II– U1 snRNP complex. Y126 of SNRNP70 is located within the RNA recognition motif (RRM) and at the interaction interface with RPB2. The crystal structure was obtained from the Protein Data Bank (PDB: 7B0Y). Bar plots on the right show the phospho-probing efficiency of tyrosine residues in RNA-preserved (RNase(-)) and RNA-digested (RNase(+)) samples. Error bars represent standard deviations. Asterisks indicate statistical significance according to a two-tailed Welch’s t-test (**; p < 0.01).

**Figure S8.**
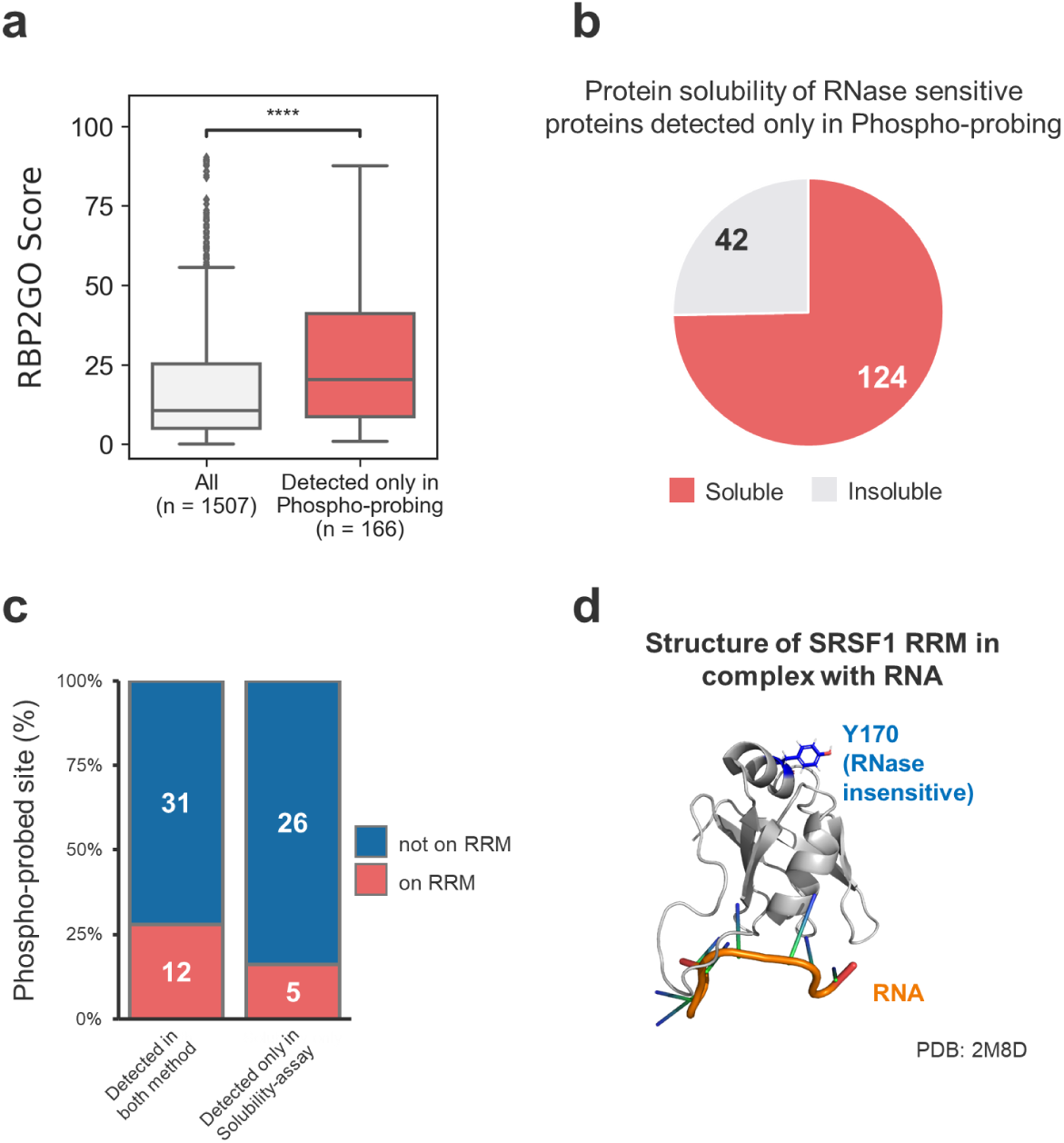
Characteristics of RNase-sensitive proteins detected by phospho-probing and solubility assay. (a) Boxplot showing the RBP2GO scores, which indicate the confidence level of RNA-binding protein annotation, for RNase-sensitive proteins uniquely identified by phospho-probing. The box spans the interquartile range; whiskers represent the 5th and 95th percentiles. The thick horizontal line indicates the median. Asterisks denote statistical significance by the Mann–Whitney U test (****; p < 0.0001). (b) Solubility classification of proteins uniquely identified by phospho-probing, based on the criteria reported by Sindhuja *et al*.. (c) Classification of phospho-probed sites among proteins detected by both solubility assay and phospho-probing, and those uniquely identified by the solubility assay. (d) Structural mapping of RNase-insensitive tyrosine residue Y170 (indicated in blue) onto the solution structure of the SRSF1 RRM–RNA complex (PDB: 2M8D). SRSF1 was identified as RNase-sensitive by the solubility assay, but not by phospho-probing.

